# White-matter degradation and dynamical compensation support age-related functional alterations in human brain

**DOI:** 10.1101/2021.12.30.474565

**Authors:** Spase Petkoski, Petra Ritter, Viktor K. Jirsa

## Abstract

Structural connectivity of the brain at different ages is analyzed using diffusion-weighted Magnetic Resonance Imaging (MRI) data. The largest decrease of the number and average length of streamlines is found for the long inter-hemispheric links, with the strongest impact for frontal regions. From the BOLD functional MRI (fMRI) time series we identify age-related changes of dynamic functional connectivity (dFC) and spatial covariation features of the FC links captured by metaconnectivity (MC). They indicate more constant dFC, but wider range and variance of MC. Finally we applied computational whole-brain network model based on oscillators, which mechanistically expresses the impact of the spatio-temporal structure of the brain (weights and the delays) to the dynamics. With this we tested several hypothesis, which revealed that the spatio-temporal reorganization of the brain with ageing, supports the observed functional fingerprints only if the model accounts for: (i) compensation of the individual brains for the overall loss of structural connectivity, and (ii) decrease of propagation velocity due to the loss of myelination. We also show that having these two conditions, it is sufficient to decompose the time-delays as bimodal distribution that only distinguishes between intra- and inter-hemispheric delays, and that the same working point also captures the static FC the best.

## 1. Introduction

Combining network dynamics approaches with graph theory metrics is a leading paradigm for studying the brain. The former is motivated by the observations that the brain architecture shapes its neurophysiological activity (Sporns et al., 2004; Vincent et al., 2007; Wang et al., 2013). As a consequence in the study of its large-scale dynamics, the brain is typically represented as a network of anatomically interacting regions each constrained by inherent dynamics (Sanz-Leon et al., 2015). Using this paradigm, a number of modeling studies have demonstrated that independently of the size of the brain regions and their underlying dynamics, neuroanatomical constraints of the human brain shape and drive its functionality during resting healthy state (Kringelbach et al., 2015; Deco et al., 2009, 2011; Schirner et al., 2018; Courtiol et al., 2020), or during pathologies such as epilepsy (Jirsa et al., 2017), stroke (Allegra Mascaro et al., 2020), or Alzheimer’s disease (Stefanovski et al., 2019).

Development of the network neuroscience (Bassett and Sporns, 2017; Bullmore and Sporns, 2009) has been possible due to advances of non-invasive structural (Johansen-Berg and Rushworth, 2009; Hagmann et al., 2010) and functional (Logothetis et al., 2001) brain imaging. The former allows application of graph theory metrics to describe the biologically realistic connectivity, the so-called connectome (Ghosh et al., 2008; Sporns et al.,2005). On the functional imaging side, until recently the focus of research was on the functional connectivity (FC) that calculates co-activation or information flow between fluctuations in the blood oxygenation-level dependent (BOLD) fMRI of distant brain regions (Friston, 2011). It is now established that non-stationarity in FC reveals a rich structure characterized by rapid transitions between few discrete FC states, which is captured by the so called dynamic Functional Connectivity (Hansen et al., 2015; Betzel et al., 2016; Calhoun et al., 2014; Allen et al., 2014). Temporal variations of the resting-state have shown to be more predictive in the context of ageing, showing differences in the modularity (Viviano et al., 2017) and decrease of FC variation for inter-modular connections (Chen et al., 2017). Dynamic FC tends to slow down and becomes less complex as well as more random with increasing age (Battaglia et al., 2020), while modular slowing of dFC was associated with cognitive dysfunction (Lombardo et al., 2020).

Ageing of the brain, is well described, both structurally (Peters, 2006; Perry et al., 2015; Lim et al., 2015) and functionally (Stumme et al., 2020; Battaglia et al., 2020), but the causality between them still has not been established on personalized level, besides the attempts to statistically link some of the observed patterns between the two (Betzel et al., 2014; Zimmermann et al., 2016). As an important attempt, a possible link was shown between the decrease of complexity of the brain function in ageing and its structural changes represented as long-range pruning (Nakagawa et al., 2013), while similarly a computational model was used to test the hypothesis that hub vulnerability in Alzheimer disease is due to the highest level of activity of the same regions (de Haan et al., 2012). There is a great amount of evidence in favor of the functional benefits of greater variability in neural systems (Sleimen-Malkoun et al., 2014; Garrett et al., 2011). Loss of variability with ageing is often followed with compensatory and adaptive processes (Lövdén et al., 2010), as well as dedifferentiation (Baltes and Lindenberger, 1997), which are theorized to play an important role in the cognition and dynamics (Cabeza et al., 2018). However, these have not been linked to the structural loss, which is consistently shown not to have a significant role in age-related cognitive decline (Burke and Barnes, 2006).

Anatomical evidence suggests that ageing mostly affects interhemispheric structural links that decrease with advancing age (Knott and Harr, 1997; Duffy et al., 1996; Kikuchi et al., 2000). Hemispheric networks decrease in efficiency with age (Caeyenberghs and Leemans, 2014), while topology of hub regions (Perry et al., 2015) and modular organization (Lim et al., 2015) remain largely stable despite a substantial overall decrease in the number of streamlines with age. Most studies show greatest deficit in the frontal regions of the brain, and there are regional differences in white matter hyperintensities (Peters, 2006), which most likely result from demyelination and reduce the axons transmission speed (Wen and Sachdev, 2004). Specifically, there is strong evidence that age and loss of myelin integrity reduces conduction velocity along nerve fibers (Peters, 2002). One of the possible mechanisms are myelin sheath alterations, which are strongly correlated with ageing in monkeys (Peters and Sethares, 2002). As for the direct evidence of conduction velocity reduction, in cats for example has been found that along nerve fibers in the pyramidal tracts it is decreased by 43% for old cats compared to young ones (Xi et al., 1999).

The work that we present has three main goals: (i) to characterize the spatial distribution of the decrease in structural connectivity including the lengths of white matter tracts, (ii) to identify the changes in the dynamic of the Functional Connectivity and its higher order spatial features, and (iii) to test the causality between (i) and (ii) on personalized level using a mechanistic model. A special attention for the structural analysis is put on the tract lengths that have been so far overlooked, even though together with the propagation velocity they define the time-delays due to axonal propagation, which is determinant for the oscillatory processes (Petkoski et al., 2018; Petkoski and Jirsa, 2019). Time delays and the weights compose the space-time structure of the brain that is crucial in shaping its macroscopic activity (Sanz-Leon et al., 2015; Ghosh et al., 2008; Deco et al., 2009), and taken together in the renormalized connectome (Petkoski and Jirsa, 2020), they unveil structural affinity for spectral activation patterns in the brain through graph theoretical metrics. Functional alterations are studied through the dFC and MC, which capture temporal and spatial aspects of the FC (Arbabyazd et al., 2020). For the last part, we utilize Kuramoto oscillators, which despite being overtly simple, due to their parsimonious parametrization allow for drawing specific links between network structure and the emergent synchronization patterns of the neuronal activity (Pope et al., 2021; Cabral et al., 2011; Allegra Mascaro et al., 2020).

## 2. Results

### 2.1. White matter loss

#### 2.1.1. Global changes in the SC

A known feature of the connectome of ageing brain, is the decrease of the total number of connections. Results in Fig. 1 show that this is robustly reflected in all three metrics used for the weights: raw counts, distinct connection counts, and weighted distinct connection counts (Schirner et al., 2015). Mostly affected are the weighted distinct connection counts, which are taking into account the surface of the GWI, while avoiding multiple counting of different tracks, and as such are mostly used in the analysis as a default connection weights (Battaglia et al., 2020; Schirner et al., 2018).

**Figure 1:**
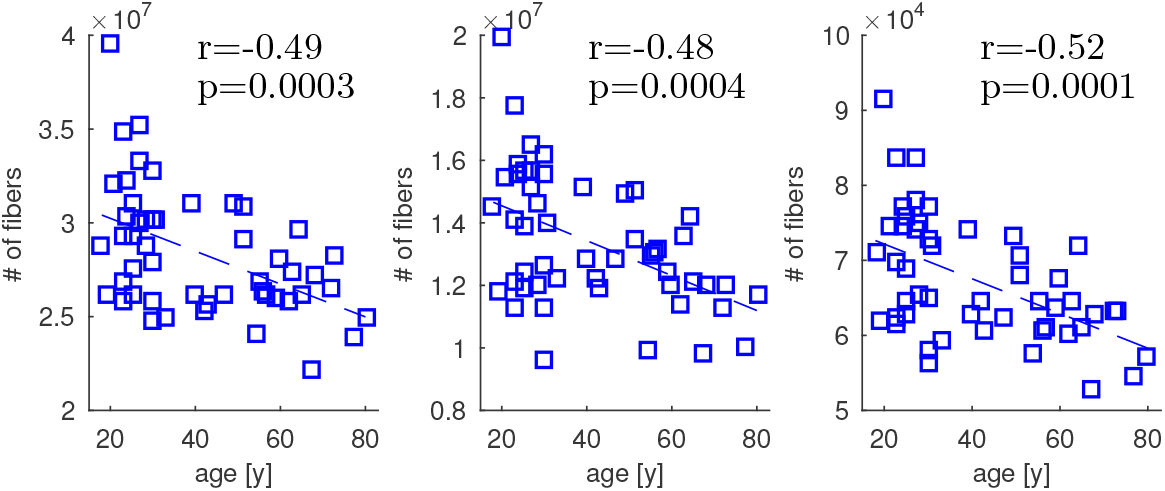
Total number of raw counts (left), distinct connection counts (middle), and weighted distinct connection counts (right) for each subject, and their correlation with age.

#### 2.1.2. Spatial changes in the SC

Next we investigated if the drop in the number of connections is spatially uniform and to what extent it is affecting the links of different lengths. We analyzed the changes of the intra- and interhemispheric links, and of the long and the short connections, where the boundary is set at 70*mm*, so that it correspond to the local minimum in the global distribution shown in the supplementary Fig. S1. These results are shown in Fig. 2, and they imply that the loss of connections is spatially and across lengths heterogeneous, with long and external links the strongest affected. For example, the number of external connections in the oldest subjects is several times smaller compared to the youngest. Since external links are longer than the internal, Fig. S1, the decreases in the external and generally longer links, Fig. 2, could be an effect of a same phenomenon targeting either longer or the external links.

**Figure 2:**
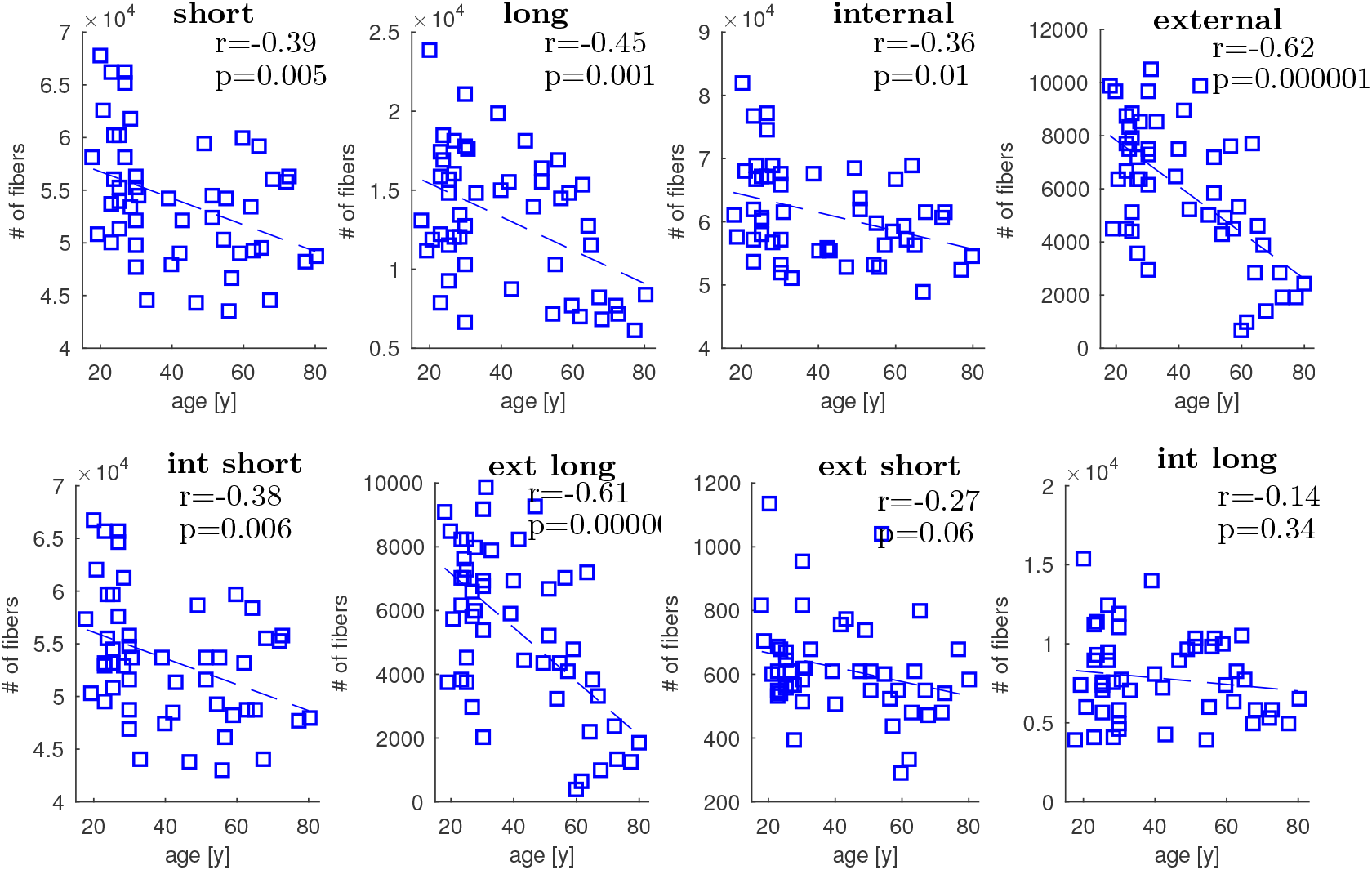
Total number of distinct connection counts, for (upper plots left to right) short, long, internal, and external links; and for (lower plots left to right) internal short, external long, external short, and internal long links.

We also checked to what extent the decrease of number of tracts impacts the average length of the remaining tracts. For each link we assume that there are as many tracts as given by the weight of the link (i.e. the counts), each of them with a length equal to the mean of all the tracts. Importantly, the overall distribution of tract lengths is quite robust across the different procedures for recovering the tracts, or if median was used instead of the mean length (see Fig. S1). Results in Fig. 3 show that the mean and median length of the tracts decrease with ageing, and that the external and long links are mostly affected. It is worth noting that the internal links have longer mean than median values, indicating that they have assymetric distribution with few very long outliers. For the external links on contrary, median and mean values are very close to each other, with short external links having much more short outliers and long external links having more long outliers that skew the means. These effects become stronger with ageing indicating that despite the significant decrease of the average length of the external links, the very long and very short external links are less affected.

**Figure 3:**
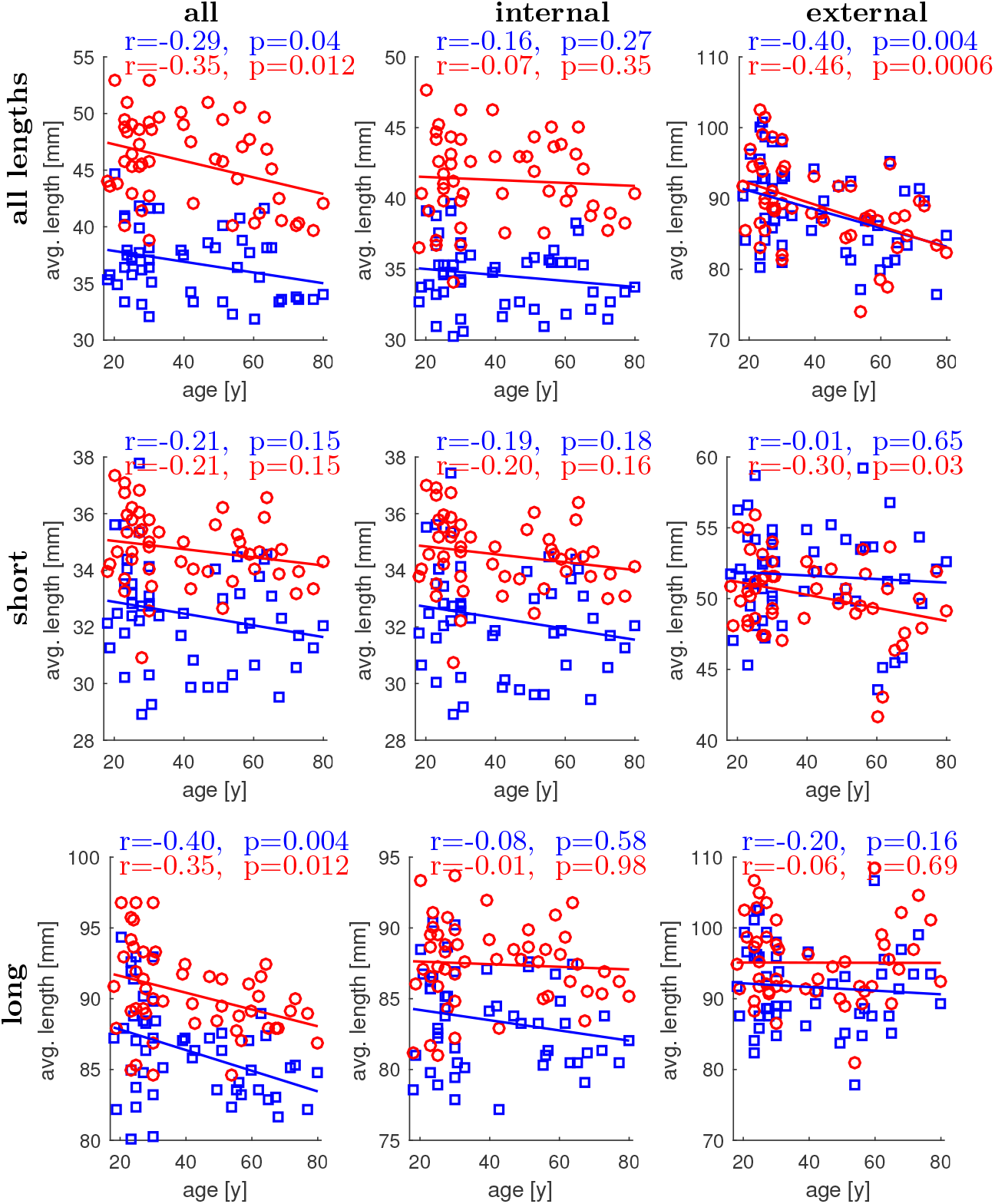
Average (blue for median and red for mean) track lengths for the whole brain (left column), internal links (middle column), and external links (right column) and for links of all lengths (top row), short links only (middle row) and long links only (bottom row).

#### 2.1.3. Lobe-specific changes

To get a better insight into the spatial extent of the white matter loss, the total number of tracts and their length was analyzed taking into account which lobes of the brain they connect. The fontal lobe is the most affected by the decrease of fibre counts, Tab. 1, and within-lobe fibers have generally larger loss than those between. Regions of the cingulate preserve the internal tracts, while their connections to the frontal, occipital and parietal lobe are significantly reduced. Besides the frontal, the number of internal tracts in the occipital and parietal lobe also experience strong negative relationship with ageing. As for the between-lobes connections, those between temporal and occipital lobes are the most decreased with age.

**Table 1:** Correlation coefficients with age and their p-values (in brackets) for the number of tracts in and between different lobes. Statistically significant values (*p* < 0.05) are bold, values with *p* < 0.005, *p* < 0.0005 and *p* < 0.00005 are indicated with one, two and three asterisks respectively.

|  | Temporal | Cingulate | Frontal | Occipital | Parietal |
| --- | --- | --- | --- | --- | --- |
| Temporal | -0.25<br>(8.6e-2) | -0.16<br>(2.5e-1) | 0.02<br>(8.9e-1) | **-0.55\*\*\***<br>(3.9e-5) | **-0.43\***<br>(1.8e-3) |
| Cingulate | -0.16<br>(2.5e-1) | 0.21<br>(1.4e-1) | **-0.53\*\***<br>(9.0e-5) | **-0.30**<br>(3.1e-2) | **-0.47\***<br>(5.3e-4) |
| Frontal | 0.02<br>(8.9e-1) | **-0.53\*\***<br>(9.0e-5) | **-0.58\*\*\***<br>(8.4e-6) | 0.03<br>(8.3e-1) | -0.01<br>(9.2e-1) |
| Occipital | **-0.55\*\*\***<br>(3.9e-5) | **-0.30**<br>(3.1e-2) | 0.03<br>(8.3e-1) | **-0.53\*\***<br>(8.1e-5) | -0.24<br>(9.5e-2) |
| Parietal | **-0.43\***<br>(1.8e-3) | **-0.47\***<br>(5.3e-4) | -0.01<br>(9.2e-1) | -0.24<br>(9.5e-2) | **-0.48\*\***<br>(4.8e-4) |

We also analyzed whether the observed effects are identically distributed among the longer and the shorter part of the links (see Tab. S1 in the Suplementary Material), where the division is set at 70*mm*. Longer tracts are much more affected than the short only for the links within parietal and between the parietal and occipital lobe. Shorter tracts on the other hand are more contributing to loss of connectivity for the links connecting the cingulate with the frontal and parietal regions, as well as those between parietal and temporal lobes. For the other significant changes from Tab. 1, the effect is mostly equally affecting longer and shorter tracts.

Another aspect of the white matter loss is shown in Tab. 2, where the decrease of number of tracts on the level of lobes is analyzed between and within hemispheres. The strongest effects from within the lobes, Tab. 1, are shown to be mainly due to the loss of interhemispheric links, for the case of the frontal and parietal lobe, while for the occipital lobe the loss is quite homogeneous. As for the connectivity between the lobes, the decrease within the hemisphere seems to be stronger. In this case the results are the same as for the short links, with parietal links to temporal and to the cingulate regions mostly affected, together with the occipital to temporal connections. On the other hand, frontal to parietal links are mostly affected by the interhemispheric connectivity loss between the different lobes.

**Table 2:** Correlation coefficients with age and p-values (in brackets) for the number of tracts in and between different lobes. Statistically significant values (*p* < 0.05) are bold, values with *p* < 0.005 and *p* < 0.0005 are indicated with one asterisks and two asterisks respectively.

| Lobes |  | Temporal |  | Cingulate |  | Frontal |  | Occipital |  | Parietal |  |
| --- | --- | --- | --- | --- | --- | --- | --- | --- | --- | --- | --- |
|  |  | left | right | left | right | left | right | left | right | left | right |
| Temporal | left | -0.13<br>(3.7e-1) | -0.05<br>(7.6e-1) | 0.18<br>(2.0e-1) | 0.07<br>(6.1e-2) | -0.06<br>(6.9e-1) | -0.16<br>(2.8e-1) | **-0.54\*\*\***<br>(5.0e-5) | -0.15<br>(3.1e-1) | **-0.35**<br>(1.3e-2) | -0.26<br>(6.7e-2) |
|  | right |  | **-0.32**<br>(2.4e-2) | 0.24<br>(8.8e-2) | 0.17<br>(2.3e-1) | -0.21<br>(1.4e-1) | 0.11<br>(4.6e-1) | -0.19<br>(1.9e-1) | **-0.38**<br>(7.2e-3) | **-0.36**<br>(1.1e-2) | **-0.39**<br>(5.3e-3) |
| Cingulate | left |  |  | 0.26<br>(7.2e-2) | 0.05<br>(7.2e-1) | **-0.43\***<br>(1.8e-3) | **-0.39**<br>(5.6e-3) | **-0.38**<br>(6.4e-3) | **-0.35**<br>(1.3e-2) | **-0.45\*\***<br>(9.1e-4) | -0.12<br>(4.2e-1) |
|  | right |  |  |  | 0.15<br>(3.1e-1) | **-0.33**<br>(2.0e-2) | **-0.45\*\***<br>(1.2e-3) | **-0.29**<br>(3.8e-2) | 0.01<br>(9.6e-1) | -0.15<br>(3.0e-1) | **-0.38**<br>(7.3e-3) |
| Frontal | left |  |  |  |  | -0.22<br>(1.2e-1) | **-0.54\*\*\***<br>(5.3e-5) | -0.04<br>(8.0e-1) | 0.03<br>(8.5e-1) | -0.04<br>(7.9e-1) | **-0.47\*\*\***<br>(4.9e-4) |
|  | right |  |  |  |  |  | -0.23<br>(1.0e-1) | -0.23<br>(1.1e-1) | 0.06<br>(6.6e-1) | **-0.38**<br>(6.0e-3) | 0.13<br>(3.8e-1) |
| Occipital | left |  |  |  |  |  |  | **-0.40\***<br>(3.7e-1) | **-0.41\***<br>(2.9e-3) | 0.02<br>(9.1e-1) | 0.21<br>(1.5e-1) |
|  | right |  |  |  |  |  |  |  | **0.51\*\*\***<br>(1.5e-4) | **-0.31**<br>(3.0e-2) | **-0.31**<br>(3.0e-2) |
| Parietal | left |  |  |  |  |  |  |  |  | -0.03<br>(8.3e-1) | **-0.56\*\*\***<br>(2.5e-5) |
|  | right |  |  |  |  |  |  |  |  |  | -0.16<br>(2.7e-1) |

Besides region-specific change in the total number of tracts, we also analyzed the relationship between age and mean of the inter and intra lobe tract lengths, since the overall length of tracts is also affected by ageing, Fig. 3. Comparing the obtained trends in Tab. 3, with those for the number of tracts in Tab. 1, we point to several features of the lobe-specific connectivity reorganization with ageing. Tracts within the frontal lobe are again influenced the strongest, meaning that not just their number is mostly reduced compared to the other lobes, but their length is mostly decreased. The same is the case with the links within the parietal and between the parietal lobe and cingulate. Moreover, separate analysis for the long versus the short portions of the tracts (see Tabs. S1 - S2 in the Supplementary Material) reveals that for the within frontal and within parietal tracts, the reduction in length is due to the strong decrease of the longer tracts, despites the preserved length by the remaining long links. As for the parietal-cingulate links, their loss mainly affects the shorter ones, which nevertheless become even shorter. It is interesting to note that temporal-cingulate links are the only one with significant increase of their average lengths, while their overall count also seems to be increasing, though not significantly.

**Table 3:** Correlation coefficients with age and their p-values (in brackets) for the mean length of tracts in and between different lobes. Statistically significant values (*p* < 0.05) are bold, values with *p* < 0.005 and *p* < 0.0005 are indicated with one asterisks and two asterisks respectively.

| Lobes | Temporal | Cingulate | Frontal | Occipital | Parietal |
| --- | --- | --- | --- | --- | --- |
| Temporal | -0.24<br>(9.9e-2) | **0.30**<br>**(3.6e-2)** | -0.09<br>(5.5e-1) | -0.11<br>(4.3e-1) | 0.04<br>(8.0e-1) |
| Cingulate | **0.30**<br>**(3.6e-2)** | 0.08<br>(5.7e-1) | -0.14<br>(3.3e-1) | -0.16<br>(2.7e-1) | **-0.40\***<br>**(4.5e-3)** |
| Frontal | -0.09<br>(5.5e-1) | -0.14<br>(3.3e-1) | **-0.50\*\***<br>**(2.0e-4)** | -0.04<br>(7.8e-1) | 0.20<br>(1.6e-1) |
| Occipital | -0.11<br>(4.3e-1) | -0.16<br>(2.7e-1) | -0.04<br>(7.8e-1) | -0.28<br>(5.2e-2) | **-0.30**<br>**(3.3e-2)** |
| Parietal | 0.04<br>(8.0e-1) | **-0.40\***<br>**(4.5e-3)** | 0.20<br>(1.6e-1) | **-0.30**<br>**(3.3e-2)** | **-0.45\***<br>**(1.0e-3)** |

The reduction of the number of tracts is not necessary related to the reduction of their length. For example, the tracts connecting the temporal with the occipital and parietal regions preserve their mean length beside the strong decrease of their total count. Similarly, although the total number of tracts between occipital and the parietal lobe is not significantly reduced, there is a significant reduction in long tracts, Tab. S1, which then leads to reduced mean length.

Regarding the changes of the average length of the links between the lobes depending on the hemisphere, the effect is generally smaller, Tab. S3. The strongest decrease is observed for the interhemispheric links within the frontal lobe, and frontal - cingulate. Interestingly, the length of interhemispheric frontal-parietal tracts are not particularly impacted besides the strong reduction in the overall connectivity Tab. 2. Similar patterns were found for the interhemispheric parietal, and for all the within occipital links.

#### 2.1.4. Clustering

Possible changes in the clustering and modularity of the SC were investigated using Louvain modularity (Blondel et al., 2008) for weighted undirected links from the Brain Connectivity Toolbox (Rubinov and Sporns, 2010). The optimal number of clusters was set at 2, 3, 4 and 5 and even though the modularity decreased in all the studied cases, Fig. 4, these were not statistically significant. For all but one subject the modularity decreases by increasing the number of partitions. Hence for 49 of the subjects statistically significant highest modularity is achieved for 2 partitions. For 32 out of 50 subjects even without setting the initial division on hemispheres, it was found as optimal one (with highest modularity coefficient) in each of the 100 runs of the algorithm. In only three of the rest, the same resolution parameters for the hemispheric division did not yield higher modularity.

**Figure 4:**
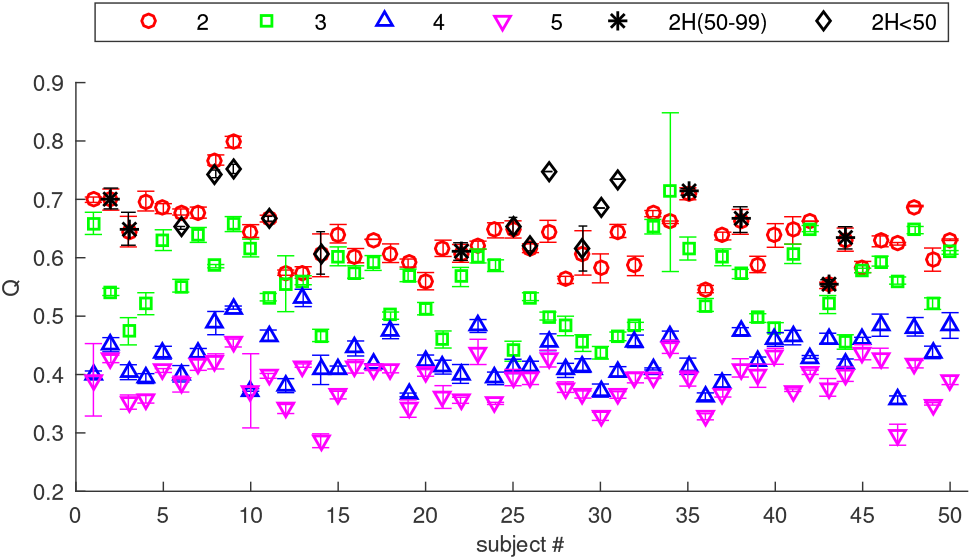
Mean modularity coefficient and their errorbars for the connectome of each subject for the most optimal partitions with 2, 3, 4 and 5 clusters, and for preset hemispheric division in the cases when not all of the 100 realizations of bicluster partitions correspond to the hemispheric division.

Similar results were observed for the modularity of weak, strong, short and long tracts, Fig. S2. As expected strong links are mostly influencing the modularity, so the results for the clustering of strong links are very similar to the clustering of the full connectome. Short links-only networks yield optimal hemispheric division for each of the subjects, whilst weak links give optimal modularity with 2 clusters for every subject, with hemispheric division being the most optimal for 45 of them. Long links on the other hand do not exclusively produce significantly smaller modularity for larger partitions, as it would have been expected from the distribution of the track lengths, Fig. S1 and also (Petkoski et al., 2016, 2018), which show that most of the interhemispheric tracts are long, compared to the median. However, due to the missrepresentation of the interhemispheric tracts with DTI (Finger et al., 2016; Jones et al., 2013; Reveley et al., 2015; Zalesky et al., 2016), the intrahemispheric long links are still of a similar size as the interhemispheric links, thus still resulting in high intrahemispheric connectivity, and hence high modularity for hemispheric division.

Taken together with the other results in this Section this suggests that the spatio-temporal structural architecture can be indeed divided into two modules corresponding to the hemispheres. Correspondingly, most of the network dynamics measures are expected to distinguish between within- and between-hemispheres large-scale dynamics.

### 2.2. Functional reorganization

First we analyzed static FC, Fig. 5 (A), for the internal and external links of each of the lobes (frontal, temporal, parietal, occipital, and cingulate). They all decrease with age, but none of the changes is statistically significant (see Tab. S4 top). Similarly, FC decreases for the internal and the external links of each of the resting state networks (Default Mode Network, Visual, Sensory Motor, Frontal Parietal, Executive Control and Auditory), but none of this yields statistical significance (see Tab. S4 bottom).

**Figure 5:**
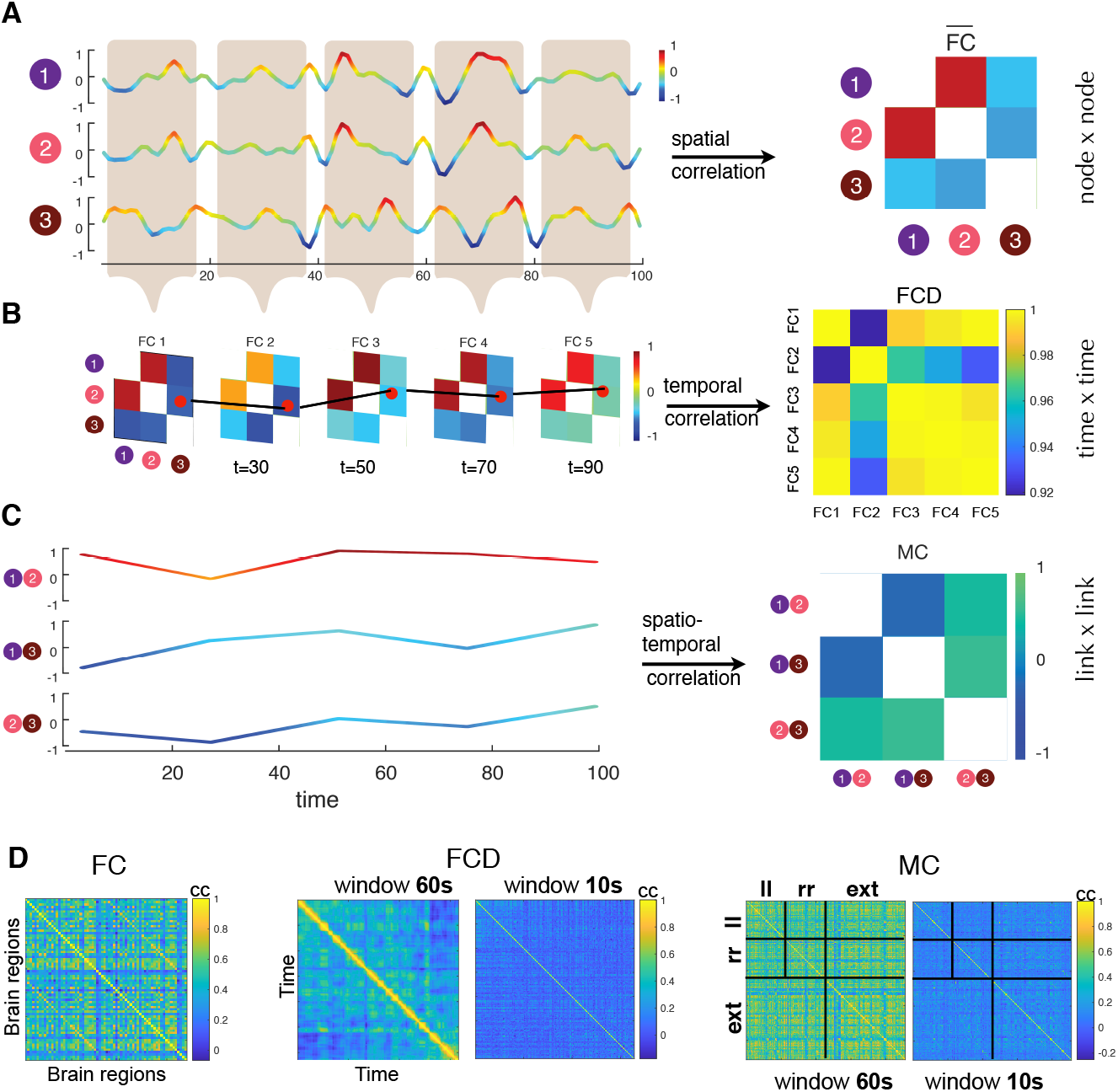
From static to dynamic Functional Connectivity. (A) Traditionally, correlations between neural activity time-series of *N* different brain region nodes are averaged over long times and compiled into the entries *FC_ij_* of a static *NxN* FC matrix (right). (B) Sliding windows of a shorter temporal duration, it is possible to estimate a stream of time-resolved *FC*(*t*) networks, so-called dFC stream. The degree of similarity (inter-matrix correlation) between *FC*(*t*) networks observed at different times is then represented into a dFC matrix. (C) Alternatively, one can consider each individual FC link as a dynamic variable *FC_ij_* attached to the graph edge between two regions *i* and *j*. Generalizing the construction of the FC matrix in panel (A), we can thus extract a *N*(*N* − 1)*xN*(*N* − 1) matrix of covariance between the time-courses of different *FC_ij_* links, giving rise to the MC matrix. (see Arbabyazd et al. (2020) for more details). (D) Example of empirical FC, FCD and MC for the first subjects, with the latter 2 calculated for window sizes of 60 and 10 seconds, and overlap of 58 and 8 respectively. Links in the MC matrix rows and columns are ordered starting from internal left hemispheric, internal right hemispheric and external (inter-hemispheric).

To go beyond the static data features, we analyzed the alterations with age in the dynamics of the FC, as captured by dFC and MC, (Arbabyazd et al., 2020), Fig. 5 (B). We focused on the dFC walk paradigm, as introduced by Battaglia et al. (2020). In short, relatively small variations of FC from one observation time to the next result in shorter flight lengths and more extensive network reconfigurations than in larger flight lengths. We characterize this temporal evolution of the FC by the mean of the dFC. Smaller mean of dFC corresponds to more fluid brain dynamics, which quicker reconfigures its dynamics.

For the spatial aspects of the dFC, we analyzed Metaconnectivity (MC) (Lombardo et al., 2020). It captures correlations between the links, Fig. 5 (C), which we grouped in different modules depending on whether they are inter- or intrahemispheric, where the latter are further divided by the hemispheres. Thus, we analyzed dynamics associated with five spatially separate modules: left-left to left-left, left-left to right-right, right-right to right-right, left-right to left-right, and intrahemispheric (left-left and right-right) to interhemispheric (left-right), Fig. 5 (D).

Fluidity of the dFC showed a significant decrease with ageing as captured by the increase of the mean non-overlapping off-diagonal dFC (further simply referred as mean dFC), and hence decrease of the mean dFC velocity, as already reported (Battaglia et al., 2020). Interestingly, the relationship is increasing and becoming more significant for decreasing length of the sliding window, Fig. 6. Similar trends are also observed for the median and the mode of the dFC, and only for the short sliding windows these are accompanied by increase of the variance and kurtosis od dFC, Fig. S3.

**Figure 6:**
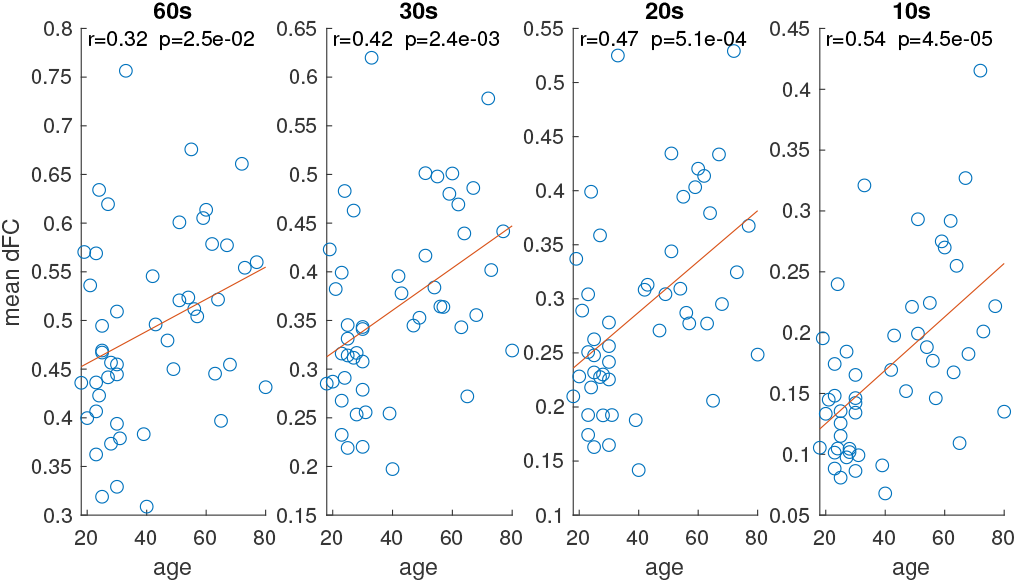
Scatter plots and the correlation of the mean dFC as a metric of dynamical reconfigurations, against the age of the subjects for different sizes of the sliding window.

Even more significant changes were observed for the higher order interactions of dFC, as captured by the MC, Fig. 7. These show very significant increase of the range and standard deviation of the values, which point to increased variations of spatial covariations. The effects were also systematically more pronounced for smaller windows. Here we were able to analyze the spatial component of the dynamical connectivity, revealing that the trends are generally preserved among all pairs of links. Nevertheless, interactions involving interhemispheric links, are slightly stronger affected, followed by the MC between the intrahemispheric links, and between the inter and intrahemispheric links.

**Figure 7:**
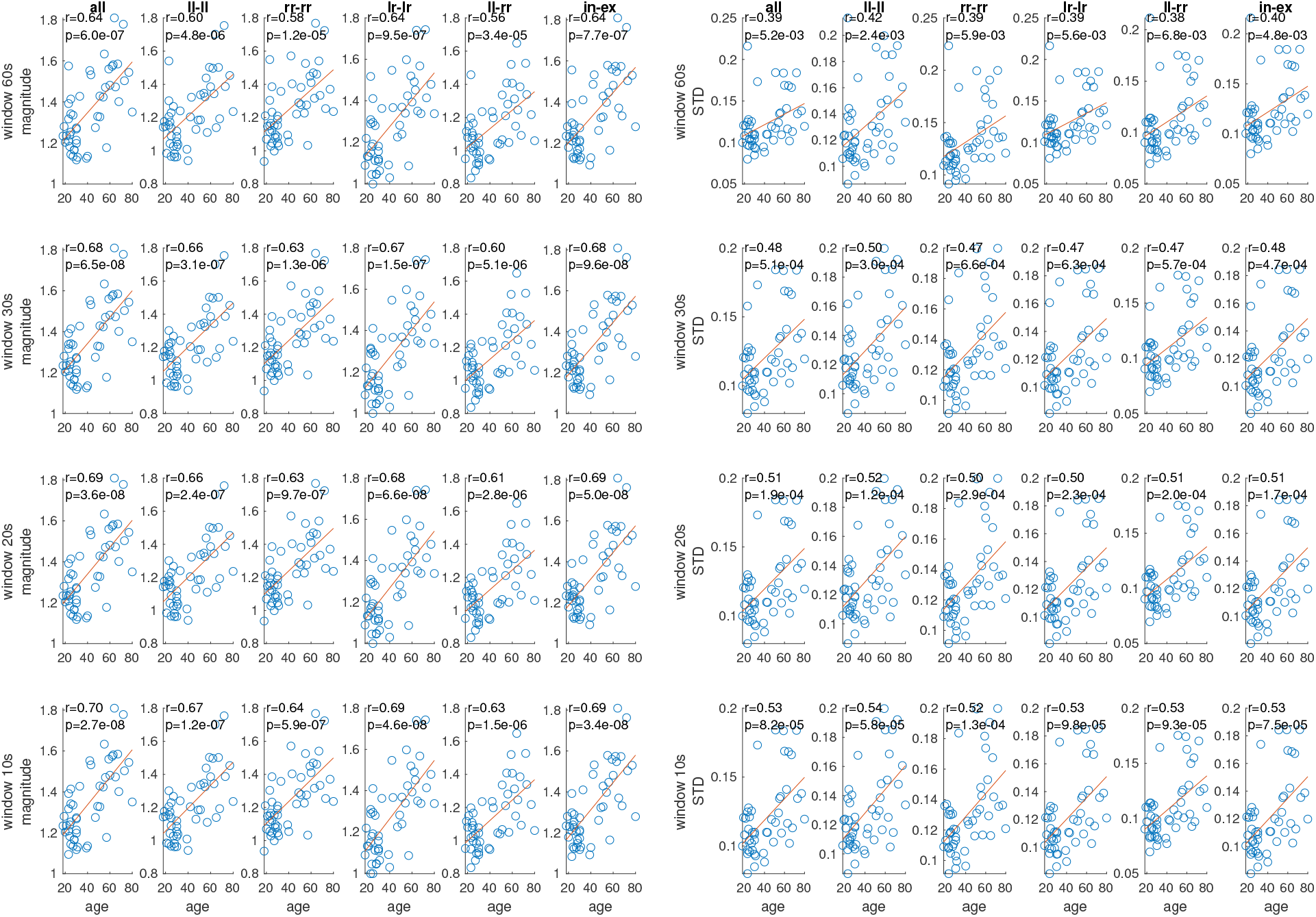
Correlation with age of magnitude (left) and standard deviation (right) of different spatial components of the empirical MC for different sizes of the sliding window.

The very strong increase of variance and magnitude of MC is followed by more modest, but still generally significant decrease in its kurtosis and mean values Fig. S4. Same as before, covariations between interhemispheric and between inter- and intra-hemispheric links are the strongest affected.

### 2.3. Brain Network Model for causal impact of SC over dFC

We constructed a brain network model to mechanistically analyze the effects of the ageing-related SC reorganization, to the neuronal activity. Even though the latter operates at EEG frequency bands, we also simulated the impact to the BOLD time-series using Balloon-Windkessel model (Friston et al., 2003) of TVB (Sanz-Leon et al., 2015), in order to be able to apply the same analysis as for the empirical measurements. We built the BNM using Kuramoto oscillators(Kuramoto, 1984) with explicit heterogeneous time-delays (Petkoski et al., 2018). Beside its simplicity and representing highly idealized system, the model can nonetheless exhibit rather non-trivial collective dynamics that could map different states of the brain (Breakspear et al., 2010;Cabral et al., 2011; Petkoski et al., 2018). In its current implementation, its simplicity allows it to only account for the impact of the spatio-temporal structure of the brain to the emergent dynamics.

Having fixed noise and natural frequencies, and personalized space-time structure, we performed parametric exploration of the global coupling and conduction velocity, as the only free parameters of the model. This allowed us to test several hypothesis. For the global coupling, we explored two strategies: (i) constant scaling for each subject, which implies a very strong impact of the connectivity loss to the dynamics, and (ii) subject-specific scaling proportional to the mean strength of the weights, which implies compensatory mechanisms for each subject. The first strategy proportionally accounts for the connectivity loss and it leads to very different dynamics between subjects, because the mean values of the weights, which mainly constrain the network dynamics (Arenas et al., 2008; Rodrigues et al., 2016) differ between subjects by almost 50% (the highest was 38.7 and lowest 22.2, with correlation with ageing of −0.2559 and p=0.0729).

The second approach decreases the influence of the SC, and practically compensate its loss by bringing the dynamical working point across subjects in the same range. However, this might be advantageous as well, since all the subjects are supposed to be healthy and in a same state. Thus it is not expected large-scale dynamical properties between them to deviate too strong, as also visible in the empirical data. Moreover, the effect of subject-specific spatio-temporal structure is still fully shaping the observed dynamics.

For the conduction velocity, we also adopted two strategies: (i) fixed velocity for each subject that does not explicitly takes into account the effects of demyelination with ageing, which are not captured by the connectome (although they are still implicitly assumed by the changes that they cause in the SC), and (ii) decreasing velocity with age, which assumes that not all of the aspects of the demyelination are fully captured by the SC through changes in the fractional anisothropy and henceforth weights, but increased propagation time at the existing links should be explicitly taken into account. In the case of age-dependent conduction velocity, it is calculated as

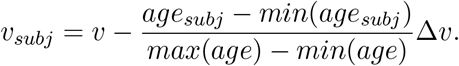

The last hypothesis that we checked with the BNM, was whether the decomposition of the time-delays into the hemispheric modes (Petkoski et al., 2016, 2018) is sufficient to capture the main data features of the dynamics at time-scales of BOLD.

Simulations for dFC reveal that there is a relatively wide parameter range for the global coupling scaled individually, and for conduction velocities linearly decreasing with age, where the average statistics of dFC is in agreement with the empirical results, Fig. 8. Significant increase for the means of dFC occurs for all 4 window lengths, for both models, with the full and with reduced delay matrices. More importantly, it is not just the same trend with age that appears statistically significant, but that is also the case for the subject specific metrics. Assuming global instead of individualized scaling for the global coupling, generally reverses the trends in the statistics of dFC, and the same is the case for constant conduction velocity.

**Figure 8:**
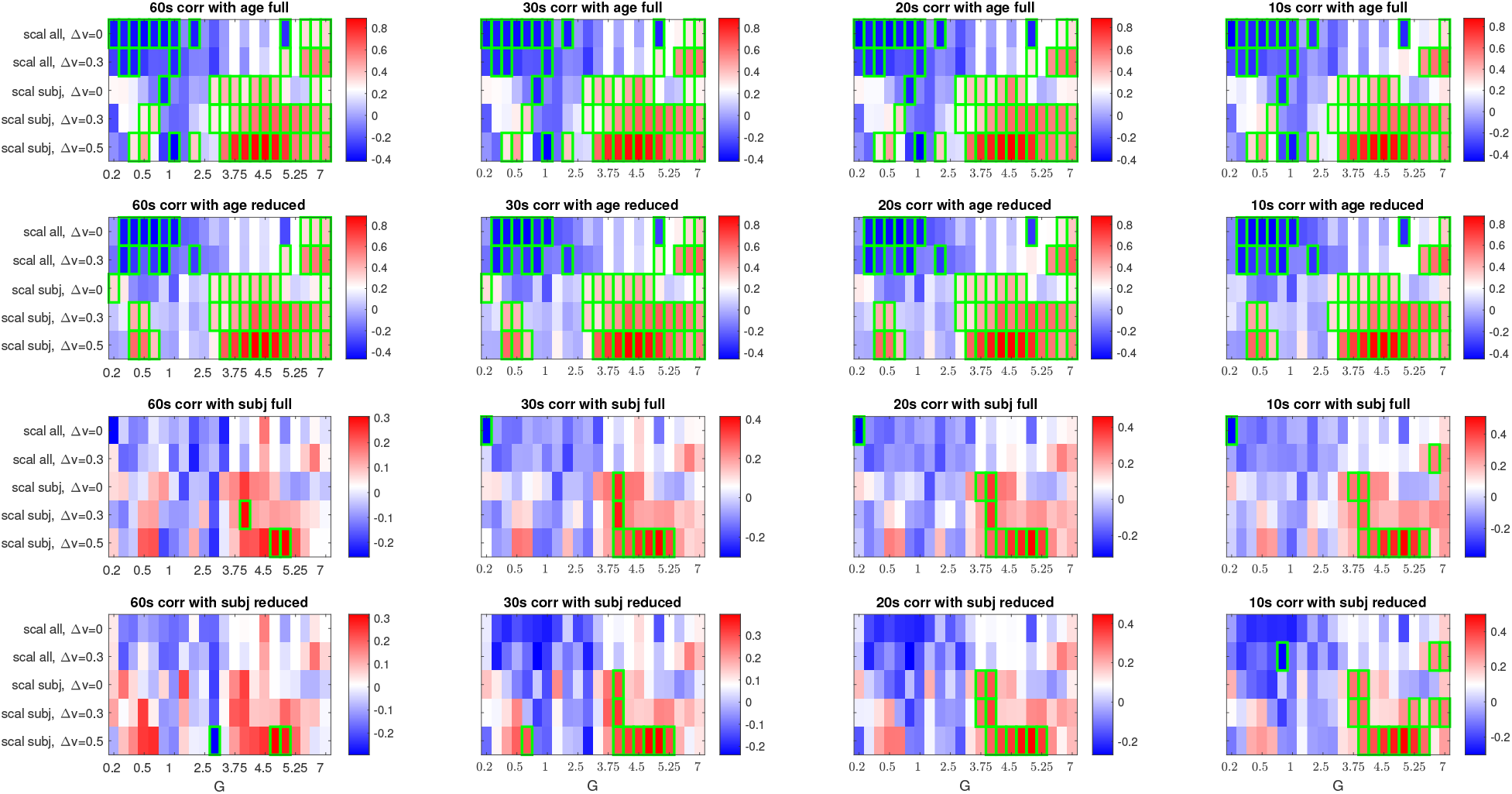
Correlation between simulated dynamical reconfigurations measures (mean of the off-diagonal FCD) with age and with the empirical results of each subject, for the full and the reduced model. The results are shown for different levels of global coupling *G*, scaling for the cohort (identical scaling for each subject indicated with *scall all* and subject specific scaling dependent on their mean connectivity indicated with *scall subj*), age dependent linear decrease of the conduction velocity Δ*v*, and window size (60*s*, 30*s*, 20*s* and 10*s*). Statistically significant correlations are indicated with green squares. Parameters: conduction velocity *v* = 3.3*m/s*, natural frequency *f* = 10*Hz*, noise intensity *D* = 1.

Same parameter sweep for the global coupling was also ran for propagation velocities of 2*m/s* and 5*m/s*, which showed qualitatively similar patterns as for 3.3*m/s*, see Fig. S3, even though less pronounced agreement with the empirical results. For the case with no delays, the results do not show any significant trends with age, nor significant correlation with empirical subject specific results.

Spatial aspect of dFC is captured by the statistics of the MC that is shown in Fig. 9. The strongest spatial patterns with age of the range and the STD of MC are well captured by both models around the same working points as for FCD. Significant correlations again appear also for the individualized dynamics, in addition to the general age-related trends. Unlike the results for FCD, here significant correlations with the ageing trends also can occur for the case of age independent propagation velocity, or global scaling, though for much fewer parameters, and never for the same working point consistently across the metrics.

**Figure 9:**
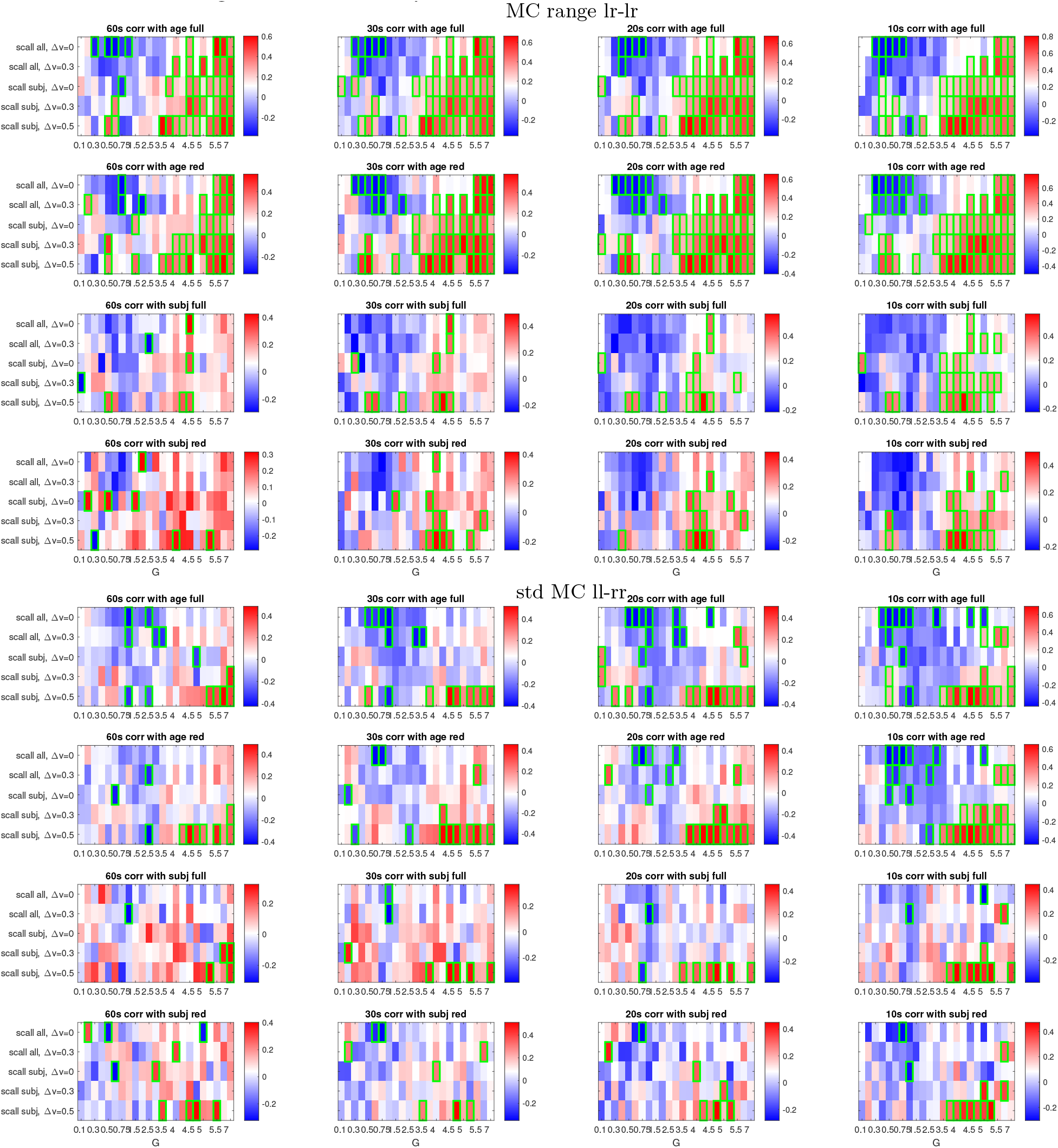
Correlation between simulated metaconnectivity measures (range in the upper plots and STD in the lower plots) with age and with the empirical results for each subject. Results are shown for different levels of global coupling *G*, scaling for the cohort (identical scaling for each subject indicated with *scall all* and subject specific scaling dependent on their mean connectivity indicated with *scall subj*), age dependent linear decrease of the conduction velocity Δ*v*, and window size (60*s*, 30*s*, 20*s* and 10*s*). The full and the reduced models are analyzed for the MC between the interhemispheric links (upper plots) and for the links in the left and in the right hemisphere (lower plots). Statistically significant correlations are indicated with green squares. Parameters: *v* = 3300*mm/s*, *D* = 1.

We performed additional exploration to characterize the dynamical working point with respect to the activity at time-scales observed in dFC. In Fig. 10 we show STD and the range of FCD values for individualized scaling of the couplings and drift of the propagation velocities, which showed the best match with the data, Fig. 8 - 9. Different window sizes show slightly different profiles for these two metrics, but interestingly, the highest predictive value of the models is observed around the values of global couplings which maximize the richness of the dFC. This becomes more obvious if the values are averaged over different window sizes (thicker red and blue lines).

**Figure 10:**
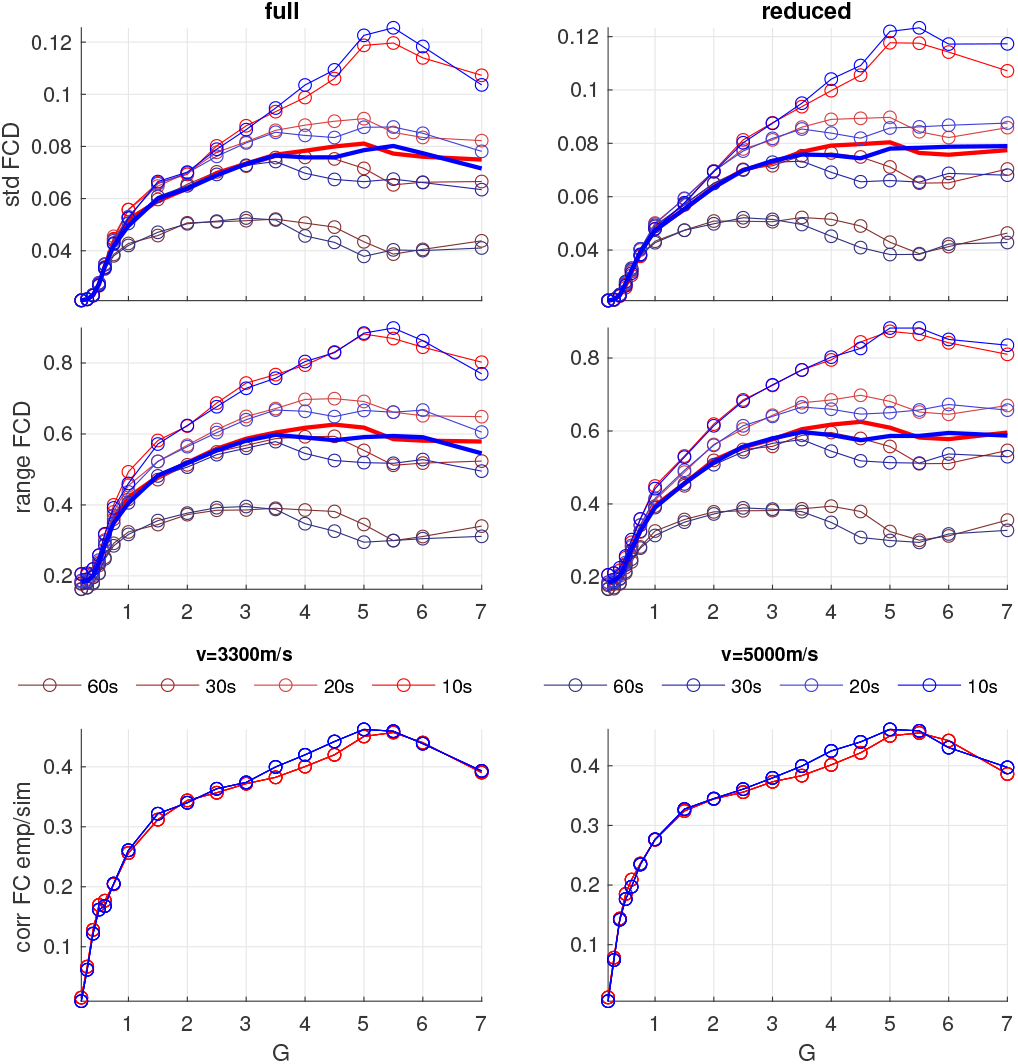
(top) STD and (middle) range of FCD of the full and reduced model for *v* = 3.3*m/s* (shades of red) and *v* = 5*m/s* (shades of blue) and for 4 different sliding windows (60*s*, 30*s*, 20*s* and 10*s*). The mean over the different sizes of the windows is also shown (thicker red and blue lines), (bottom) Mean correlation over the cohort of the individualized simulated and empirical FC. The global coupling is scaled with the mean weights for each subject, Δ*v* = 0.3, and *D* = 1.

In the same figure, we also show that these optimal dynamical range overlaps with the working point where the model reaches highest predictive value for the static FC. It is worth noting that for the fast dynamical activity, the same model has richest dynamical repertoire as captured by the variability of the Kuramoto order parameter at smaller global couplings (see Fig. S4).

## 3. Methods

### 3.1. Data acquisition

In this study we analyze anatomical and diffusion weighted images of 50 subjects acquired at Berlin Center for Advanced Imaging, Charite University Medicine, Berlin, Germany (age ranged from 18 to 80 years, mean 41.26 ± 18.36; 30 females and 20 males). 49 of these subjects were used in the study for describing the automated pipeline for constructing personalized virtual brains from multimodal neuroimaging data (Schirner et al., 2015) and their dFC were also analyzed as part of a larger cohort (Battaglia et al., 2020). The pipeline combines several state-of-the-art neuroinformatics tools to generate subject-specific cortical and subcortical parcellations, surface-tessellations, structural and functional connectomes, lead field matrices, electrical source activity estimates and region-wise aggregated BOLD fMRI time-series (Schirner et al., 2015).

### 3.2. Construction of SC networks

Desikan-Killiany atlas (Desikan et al., 2006) is used as implemented in FREESURFER (excluding the corpus callosum, but including the insular cortices of both hemispheres) leading to 68 cortical regions of interest (ROI). Since tracks are two-dimensional line objects there is no straightforward way to compute the surface area of a track-terminal. Therefore, (Schirner et al., 2015) devised a new bundling scheme in order to group tracks into distinct connections. A distinct connection is defined as a pair of grey-white matter interface (GWI) voxels for which at least one track was generated, regardless of the number of tracks found or the concrete pathway. This metric is further expanded to a second metric by another assumption that states that all distinct connections that share a common terminal voxel must also be bounded by the same bottleneck, and consequently the maximum bandwidth of that voxel must be split up among all distinct connections. Therefore, upon determining all distinct connections for a voxel, the total coupling strength of this voxel is split up over all its distinct connections in equal parts resulting in weighted distinct connections.

It follows that in this approach the maximal total coupling strength of a region is given by the area of its GWI and not by the amount of tracks that emanate from it, since this number is highly dependent on local anatomy and the characteristics of the diffusion profile. This approach is justified by the assumption that the coupling strength of a region is proportional to the size of the GWI of that region, assuming a fixed ratio of long-range connections per microcircuit volume.

Upon tractography the pipeline (Schirner et al., 2015) computes distinct connections and aggregates them for each region to generate three types of SC matrices:

- raw counts, contain track counts of all tracks that were found between each pair of regions (symmetric),
- distinct connection counts, contain only distinct connections between each pair of regions (symmetric),
- weighted distinct connection counts, in which the strength of each distinct connection is divided by the number of all distinct connections leaving the voxel (yielding asymmetric strength matrices). For the last case, symmetric SC matrix was constructed by taking the mean of the weights per each direction of the links.

Along with strengths, the pipeline outputs three different SC distances matrices that contain the mean, mode and median lengths of all tracks that were found between each pair of regions.

### 3.3. MRI acquisition

Magnetic resonance imaging (MRI) acquisition was performed on a 3T Siemens Tim Trio scanner. Every subject was scanned in a session that included a localizer sequence (3, 8 mm slices, repetition time [TR] = 20 ms, echo time [TE] = 5 ms, voxel size = 1.9 × 1.5 × 8.0mm, flip angle [FA] = 40°, field of view [FoV] = 280 mm, 192 mm matrix), a T1-weighted high-resolution image (MPRAGE sequence, 192, 1 mm sagittal slices, voxel size 1 × 1 × 1mm, TR=1940 ms, TE=2.52ms, FA = 9°, FoV = 256 mm, 256 mm matrix), a T2 weighted image (2:16 min, 48,3mm slices, voxel size 0.9 × 0.9 × 3mm, TR=2640ms, TE1=11 ms, TE2=89 ms, FoV=220 mm, 256 mm matrix), followed by diffusion weighted imaging (61, 2 mm transversal slices, voxel size = 2.3 × 2.3 × 2.3 mm, TR = 7500, TE = 86 ms, FoV = 220 mm, 96 mm matrix). Subjects were then removed from the scanner to have their EEG cap put on, and then simultaneous fMRI-EEG images were acquired in a single run (BOLD T2*weighted, 32, 3 mm transversal slices, voxel size = 3 × 3 × 3 mm, TR=1940 ms, TE=30 ms, FA= 78 °, FoV = 192 mm, 64 mm matrix). Five dummy scans were automatically discarded by the Siemens scanner.

During the scans, subjects were to remain awake and reduce head movement. Head cushions served to minimize head movement, and earplugs were provided, during the 20 min of uninterrupted scan.

### 3.4. fMRI processing

fMRI data were pre-processed following Schirner et al. (2015) and in a same manner as in Battaglia et al. (2020), using the software FEAT (fMRI Expert Analysis Tool) first-level analysis from the FMRIB (Functional MRI of the brain). Motion correction was performed using EPI field-map distortion correction, BET brain extraction, and high-pass filtering (100s) to correct for baseline signal drift, MCFLIRT to correct for head movement across the trial. As an additional correction measure, we further regressed out six FSL head motion parameters from the measured BOLD time-series. Functional data was registered to individual high-resolution T1-weighted images using linear FLIRT, followed by nonlinear FNIRT registration to Montreal Neurological Institute MNI152 standard space. Voxel-level BOLD time series were reduced to 68 different brain region-averaged time series, according to a Desikan parcellation (Desikan et al., 2006). We neither performed a slice-timing correction, smoothing, normalization of BOLD intensities to a mean, nor global regression.

### 3.5. Modularity

The optimal community structure of the connectomes was calculated using the Louvain function (Blondel et al., 2008) from the Brain Connectivity Toolbox (Rubinov and Sporns, 2010) for the weighted undirected links. For each subject the algorithm was ran starting from a value of the resolution parameter which gives twice the number of the required partitions. Then the resolution parameter was decreased by 1% at each consecutive step until a partition consisting of predefined number of communities was achieved. The procedure was repeated 100 times each, for partitions of 2, 3, 4 and 5 communities. For each of these partitions the modularity coefficient *Q* was calculated. It is a scalar value between −1 and 1 that measures the density of links inside communities as compared to links between communities (Newman and Girvan, 2004). In addition we calculated the modularity coefficient for the case when the initial community affiliation vector was set to correspond exactly to the left and right hemispheres. Here again an algorithm was ran for the same 100 resolution parameters that earlier achieved optimal bi-community structure, until it was assured that the division in 2 communities corresponds to the hemispheric division.

### 3.6. dFC metrics

Static (time-averaged) functional connectivity is calculated as Pearson correlation of the BOLD time-series, empirical or simulated, in a given time window. Evolution of FC over time is captured by dFC that here is defined the same as Functional Connectivity Dynamics (FCD) in Hansen et al. (2015). It describes the similarity between *FC*(*t*) matrices at different time windows *t_k_*, where *k* = 1 … *M*. The (*t_k_*, *t_l_*) entry of the *M* × *M* FCD matrix is provided by the Pearson correlation between the upper triangular parts of the two matrices *FC*(*t_k_*) and *FC*(*t_l_*) (Arbabyazd et al., 2020). We calculated FCD for 4 different window sizes (30, 15, 10 and 5 time points), which always move by one time point. In all the cases, by keeping the interval between two consecutive FC network time-resolved estimations constant, higher and lower correlations of the FCD can naturally be interpreted as associated to a slower or faster speed of dFC reconfiguration (Battaglia et al.,2020). Hence, a first aim to describe dFC consists of a suitable quantification and description of the distributions of FCD entries, notably, their mean, median or mode, giving a typical dFC speed, and their spread as given by their second and higher momenta, i.e. STD and kurtosis.

In addition we calculated higher order interactions between brain regions using metaconnectivity (MC) (Arbabyazd et al., 2020). Exactly as typical static FC analysis ignore time, the previously mentioned FCD analyses ignore space. However, FC reconfiguration may occur at different speeds for different sets of links (Lombardo et al., 2020). Furthermore, the fluctuations of certain FC links may covary with the fluctuation of other FC links, but in the same time be relatively independent from the fluctuation of other sets of links. Therefore, we compute a different dFC speed distribution for different sets of links, wgich constitute spatial dFC modules. MC is defined as correlation between link-wise time-series consisting of the pair-wise correlations between the given nodes at each window. Hence it represents a fourth order statistics between node’s dynamics. We have ordered the links in the MC matrices in a such a way that besides the statistics of the overall MC, we also calculate the statistics of the links within the left and within the right hemispheres, between the hemispheres, between the internal left versus right hemispheric links, and left and right versus interhemispheric each. These are all illustrated in Fig. 5, where the metaconnectivity within the 7 clusters is expected to be distinctive, as a consequence of the high hemisphiric bimodularity of the SC discussed in Section 3.5.

## 4. Model

### 4.1. Brain Network Model with Kuramoto oscillators

We build the personalized BNM comprising of 68 delay-coupled cortical brain regions, each having a dynamics captured by Kuramoto oscillators (Cabral et al., 2011; Petkoski et al., 2018;Pope et al., 2021). For the metrics of connection strengths we have chosen the weighted distinct connection counts, because they take most physiological features into account (Schirner et al., 2015). The time delays corresponding to the links are defined from the personalized lengths by setting the propagation velocity from within the physiological range for brain signals (Nunez and Srinivasan, 2006; Trebaul et al., 2018), which is usually between 1 − 10*m/s*. In particular we use 3.3*m/s*, which was shown to have a highest predictive value for supporting realistic spectral activation patterns (Petkoski and Jirsa, 2020). To confirm the generality of the results, we also use propagation velocities of 2*m/s* and 5*m/s*. For the lengths of the tracks, we use mean values, instead of median or the mode, because some links having small number of tracts so the mode cannot be reliably defined and detected, whilst the median and mean have similar values, as shown in Fig. S1.

We consider the Kuramoto model (KM) (Kuramoto, 1984) with explicit heterogeneous time-delays *τ_ij_* and coupling strengths *K_ij_*, rewritten as

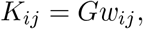

where *G* is a global coupling parameter and *w_ij_* are the normalized weights from the connectome. The model represents as a canonical model for weakly delay-coupled oscillators with long delays in comparison to the coupling strengths or natural frequencies *ω_i_* (Izhikevich, 1998; Ermentrout and Wechselberger, 2009). For symmetric, link-dependent delays, *τ_ij_* = *τ_j,i_*, phases *θ_i_* of each oscillator evolve as

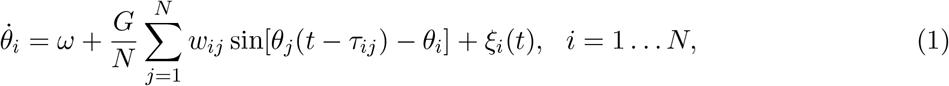

where *ω* is natural frequency of oscillators and *ξ_i_*(*t*) takes into account the contribution of different stochastic forces and are assumed to be sources of Gaussian white noise satisfying 〈*ξ_i_*(*t*)〉 = 0, 〈*ξ_i_*(*t*)*ξ_j_*(*t*)〉 = 2*Dδ_ij_δ*(*t* − *t′*)

The heterogeneity in phase models can stem from natural frequencies and/or from additive noise term, but both sources have similar influence to the observed global dynamics (Acebrón et al., 2005). The size of the connectome-derived networks is quite small, *N* = 68 cortical regions, and the connection strengths span almost across 5 orders of magnitude, hence if the natural frequencies are heterogeneous the global dynamics would be highly influenced by the particular realization of the probability density function of the natural frequencies (Petkoski et al., 2018). To avoid this we have decided to fix the natural frequencies of each node, while still introducing heterogeneity to each oscillator in a form of a independent white noises with same intensity.

For the parameters space, we explore only the global coupling *G* and the propagation velocity *v*, while keeping fixed the level of the noise *D* and the natural frequencies *f*. This is justified because the dynamics for the KM in general depends on the ratio *G/D* (Acebrón et al., 2005). Similarly the impact of the time-delays depend on their relative size compared to the natural frequencies (Petkoski et al., 2016).

### 4.2. BNM with reduced distribution of delays

The reduced model is derived from the decomposition of the space-time structure of the connectome (Petkoski et al., 2016, 2018). It assumes bimodal *δ*-distributed time-delays, with values corresponding to the mean delays of the internal and external links for each subject. The model hence read

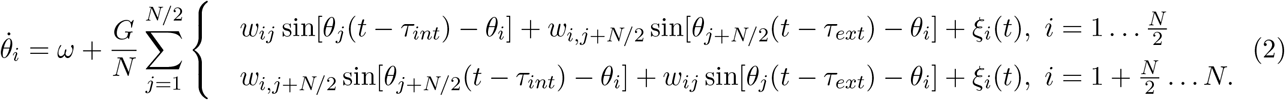

### 4.3. Simulated BOLD

Simulated neural activity was converted into simulated BOLD signals using a Balloon-Windkessel model (Friston et al., 2003) and the resulting time-series were then downsampled at 2*s*, to correspond to the empirical BOLD signals. Because of the simplicity of the model, the physical quantity whose variations underlie BOLD signal was chosen as *b_i_* = sin*θ_i_*, as it has been the case in other studies of the neuronal activity using Kuramoto oscillators (Cabral et al., 2011; Ponce-Alvarez et al., 2015; Petkoski et al., 2018; Pope et al., 2021).

## 5. Discussion

In this work we identify the significant changes in the dynamics of brain FC, as reflected in dFC and MC metrics for the dynamical reconfigurations, and we apply a BNM to causally test which aspects of the spatiotemporal reorganization of the brain structure with the ageing, could be responsible for the former dynamical alterations.

### 5.1. SC alterations with age

Age-related changes of the total number and the weights of tracts have been of interest of many studies (Perry et al., 2015; Betzel et al., 2014; Lim et al., 2015) that, nevertheless, overlooked the spatial distribution of tract lengths on which we also focus here. The most commonly observed feature of the topological organization of the white matter connectivity across the human lifespan is an inverted U shape (Zhao et al., 2015; Coupé et al., 2017), with a decrease in the global network properties, such as the network strength, starting from the third decade, as confirmed with our results for the decrease in SC. This loss of white matter connectivity is especially pronounced for the inter-hemispheric links, which is also in line with the literature (Puxeddu et al., 2020). This on the other hand also causes the average length of the remaining tracts do decrease with age, since the interhemispheric links are on average longer.

The mostly affected area by the decrease of the fibre counts was frontal lobe, Tab. 1, which was in agreement with the similar studies (Gunning-Dixon et al., 2009; Peters, 2006). Similarly strong loss is observed in the occipital lobe, but this is more equally affecting intra- and inter-hemispheric links, hence having much smaller impact on the tract lengths. It is interesting to be noted that even though we found within-lobe fibers to be more affected than those between lobes, Tab. 1, the modularity is still generally stable, which is in line with the literature (Lim et al., 2015).

In addition, many studies indirectly point to the decrease of white matter fibers in the frontal regions by showing strong negative relationship with functional anisotropy (FA) and age, that is especially prominent in the frontal lobe (Grieve et al., 2007; Bennett et al., 2010; Billiet et al., 2015;Inano et al., 2011), whereas (Sexton et al., 2014) reported a general accelerated decline in anisotropy during senescence. Most of the these works (Grieve et al., 2007; Mädler et al., 2008; Inano et al., 2011) also measured axonal and radial diffusion and related the trends in those measures to the demyelination, as indicated by the measurements of myelin water fraction. However, there are also some contradicting studies (Billiet et al., 2015), which although observed the decrease of the FA with ageing, did not find significant difference in myelin water fraction. We did not recover myelination or other related metric in this study, but instead using the BNM we found that model’s predictability is increased if a reduction of conduction velocity due to hypothesized demyelination is assumed to occur linearly with age.

### 5.2. dFC alterations with age

It is now a widely recognized concept in the study of the dynamics of the human brain network, that functional connectivity is not static, but changes its pattern over time, even during rest. The possibility of studying these dynamics through careful analysis of neuroimaging data has catalyzed substantial interest in methods that estimate time-resolved fluctuations in functional connectivity (Lurie et al., 2020). In the context of ageing, analysis of the temporal variations of the resting-state showed differences in the modularity (Viviano et al., 2017), and decrease of FC variation for the inter-network connections (Chen et al., 2017). Dynamic FC tends to slow down and becomes less complex as well as more random with increasing age (Battaglia et al., 2020), and similarly, modular slowing of dFC was associated with cognitive dysfunction induced by sleep deprivation (Lombardo et al., 2020). Cognitive performance in healthy older adults relates to spontaneous switching between states of functional connectivity during rest, as captured by FCD (Cabral et al., 2017). Also, lower metastability at slow time-scales of BOLD activity is associated with higerh age (Escrichs et al., 2020), which is in agreement with the functional benefits of greater variability in neural systems, including flexibility/adaptability, heightened dynamic range, Bayesian optimality, and multi-stability (Sleimen-Malkoun et al., 2014; Garrett et al., 2011).

Here, first we showed that several metrics derived on static FC do not any predictive value for the ageing, which is not surprising knowing that simulated FC that fits best to the same subjects empirical FC may not necessarily be the same simulated FC (Triebkorn et al., 2020). Next, we focused our analysis on the temporal aspects of the dynamics of FC and the spatial aspects of the covariations between the FC links. The former generally confirmed the findings from (Battaglia et al., 2020) about slowing down of dFC, as reflected by it’s average values. This decrease of average dFC (Rabuffo et al., 2021) seems to accelerate at shorter time-scales, as observed from dFC calculated at sliding windows of 10s. Interestingly, the variance of dFC seems less affected, and only starts to significantly increase at faster time-scales.

Age related changes are even stronger for the covariations between the FC links, as captured by different spatial components of MC. Here it is not the mean values that are mostly affected, but the variance and the range of the MC values, which increase very significantly with age. Larger trends for shorted time-scales are also visible here, though to much smaller extent. Spatially speaking, the trends are strongest for the covariations of inter-hemispheric links (even though the interhemispheric static FC is unaffected) between themselves, and with the intra-hemispheric ones. These results point for first time that taking into account the spatial aspects of the dFC or higher order interactions, could serve as a better and more robust biomarker for ageing. This is to be expected also due to the large spatial heterogeneity of the white-matter loss, which nevertheless does not result in observable changes in the respective static FC metrics. A possible reason for this is the fact that FC by definition averages out all the nonlinearities in the dynamics (Hansen et al., 2015), which is highly non-stationary (Heitmann and Breakspear, 2017; McIntosh and Jirsa, 2019).

### 5.3. Linking the structure and the function

Age-related alterations in brain structure and function have been linked to the age-related cognitive decline, though challenges still remain (Hedden et al., 2016). Sources of heterogeneity are not fully understood, but seem to be associated with different neurobiological substrates (loss of white matter tracts and demyelination), and single-cohort designs might be optimal in reducing the sources of interindividual variation that may be unrelated to age (Zuo et al., 2016). Current models indicate that structure and function are significantly correlated, but the correspondence is not perfect because function reflects complex multisynaptic interactions in structural networks (Suárez et al., 2020). This has been demonstrated also in the respect with ageing (Zimmermann et al., 2018), where SC and FC each show unique and distinct patterns of variance across subjects, and variability of alterations of functional connectivity is especially high across older adults (Stumme et al., 2020). Therefore, function cannot be directly estimated from structure, but must be inferred by mechanistic models, which can causally test the higher-order interactions (Schirner et al., 2018; Courtiol et al., 2020), and hence offer higher explanatory value compared with the data-driven methods (Jockwitz et al., 2017). It was already shown that a specific pattern of SC/FC coupling predicts age more reliably than does region wise SC, or FC decrease alone (Zimmermann et al., 2016), but causal link between the SC age-related changes and the FC reorganizations are still ambiguous. This indicates that the impact that SC has on the brain dynamics need to be investigated beyond the FC (Triebkorn et al., 2020).

Compensation (Lövdén et al., 2010) and dedifferentiation (Baltes and Lindenberger, 1997) of brain are probably among the major causes that make establishing the SC/FC link with ageing on individualized level even more difficult. Compensatory mechanisms of the global coupling in shifting the dynamical working point were already demonstrated in the case of the resting-state of epilepsy (Courtiol et al., 2020). This homeostatic compensation is one of the main theories for the ageing (Park and Reuter-Lorenz, 2009; Sleimen-Malkoun et al., 2014). Another reason could be that time-delays are often ignored, and they still cannot be retrieved on individual level, but tract lengths are used as a proxy (Sanz-Leon et al., 2015). This however cannot capture the important effects that demyelination might have on increasing the propagation delays (Sorrentino et al., 2021a) With the model that we have constructed, we try to account for both, the impact of the age-related compensation and demyelination, combined with the individualized spatio-temporal structure.

The model shows statistically significant individualized predictive value for the most significant dFC and MC data features. This is observed only if (i) dynamical compensation is assumed through the global effective coupling that is identical across subjects, by normalizing each connectome with the individualized mean connectivity, and (ii) conduction velocity is linearly decreased with ageing to account for the demyelination that is not accounted in the weights and lengths of the individualized connectome. Moreover, the results hold even if space-time structure is spatially decomposed (Petkoski et al., 2016, 2018; Petkoski and Jirsa, 2019), such that only lumped intra and interhemispheric delays are taken into account. The latter is supported by the results for the SC alterations, which indeed show that the strongest decrease with ageing is observed for the interhemispheric tract lengths. It is also worth noting that the best fit for the mean of FCD is obtained at the working point that maximizes the variability of dFC as observed through the variance and the magnitude of the (Triebkorn et al., 2020). This holds for both the original and the reduced model.

To summarize the main results from the model are quite robust and they indicate that:

1. Brain compensates for the loss of connectivity with the age by increasing the effective global coupling. Common scaling that excludes compensation leads only to negative correlations with the empirical data, regardless of the propagation velocity, or whether it is constant or decreasing with age.
2. Using identical conduction velocity across age has very small predictive value. On contrary, significantly better predictability is achieved if the propagation velocity is assumed to linearly decrease with age due to demyelination. This is the only scenario that leads to significant correlation of the simulated results not just with age, but also with the personalized FCD and MC metrics, and it holds for different propagation velocities and strengths of decrease.
3. Spatio-temporal decomposition along its hemispheric modes, has similar predictive value as the full model, meaning that inter-hemispheric SC alterations during ageing are of the crucial importance for the dynamical alterations.
4. The working point with the best correlation is at the parameters space with highest variability at slow time-scales, and the same working point also gives a reasonable agreement with the static FC.

### 5.4. Limitations of the study

The reliability of the obtained results for the impact of the structure on the emergent function during ageing could be improved by using larger cohorts, which become more available, such as 1000Brains (Caspers et al., 2014) or the Human Connectome Project Van Essen et al. (2013). For this, it would be necessary to use digital research platforms for brain science such as EBRAINS that allows integration and accessibility of those datasets together with simulation engines (Schirner et al., 2021). Similarly, parcellation-induced variation of empirical and simulated brain connectomes at group and subject levels is another issue that needs to be considered Domhof et al. (2021). Nevertheless, most of our finding, independently for the structure and function, and for their link with the brain network model, show quite significant trends from statistical viewpoint, giving us confidence in the reported statistical changes and in the mechanisms linking them, which are recovered on individualized level for a first time.

Decomposition of time-delays in the reduced model probably works because using homogeneous velocities is already introducing lots of bias for the delays. As a such, the main modes of the delays seem to contain sufficient predictive value, as demonstrated in-silico (Petkoski et al., 2018; Petkoski and Jirsa, 2019). It should be noted that obtaining individualized conduction velocity for large scale brain models is a complex issue. One could argue that taking a single value is far from reality, since the delays are known to differ by several orders of magnitude and to be directly dependent of the diameter of the axon and the presence of a myelin sheath. Thus, action potential velocity can be generally between 0.1 m/s in unmyelinated axons and 100 m/s in large myelinated axons, and it is a direct function of the diameter of the axon and the presence of a myelin sheath (Waxman, 1980). Besides, the presence of axonal irregularities and repetitive stimulation or activation of specific ion channels also reduce the conduction velocity (Debanne, 2004). In that sense, (Caminiti et al.,2013) found that the spectrum of tract lengths obtained with MRI closely matches that estimated from histological reconstruction of axons labeled with an anterogradely transported tracer. They also measured conduction velocity of myelinated axons in human brain were between 6-10 m/s. However, it has also been reported that similar overall population of myelinated to non-myelinated axons can be found in corpus callosum of different species (Olivares et al., 2001), with the latter having generally velocity <1m/s. All these indicates that better estimates of the time-delays (Trebaul et al., 2018), especially if they are personalized (Sorrentino et al., 2021b) are expected to improve the results of any large-scale BNM.

## 6. Acknowledgements

This research was supported by the European Unions Horizon 2020 research and innovation programme under grant agreement No. 945539 (SGA3) Human Brain Project and by grant agreement No. 826421 Virtual Brain Cloud. In addition, PR acknowledges support by ERC Consolidator 683049; German Research Foundation SFB 1436 (project ID 425899996); SFB 1315 (project ID 327654276); SFB 936 (project ID 178316478), SFB-TRR 295 (project ID 424778381); SPP Computational Connectomics RI 2073/6-1, RI 2073/10-2, RI 2073/9-1; Berlin Institute of Health & Foundation Charit, Johanna Quandt Excellence Initiative.

## 7. Suplementary Material

**Figure S1:**
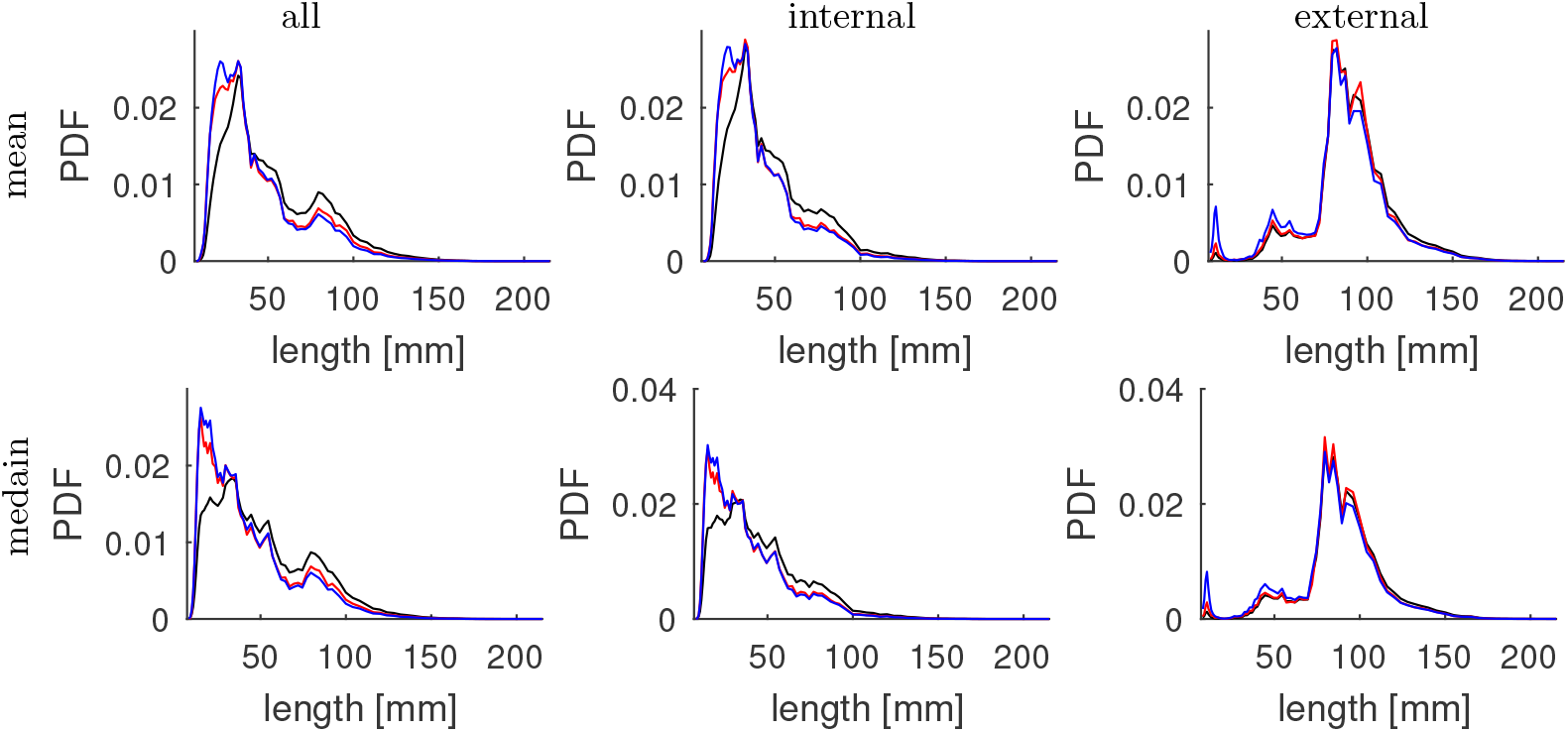
Probability Density Function (PDF) of global (left column), internal (middle column) and external (right column) weighted lengths of the links, for distinct connection counts (black), weighted distinct connection counts (red) and raw counts (blue) of the tracks. For each link, the length is calculated as a mean (upper row) and median (middle row) length of all the tracts in that link.

**Figure S2:**
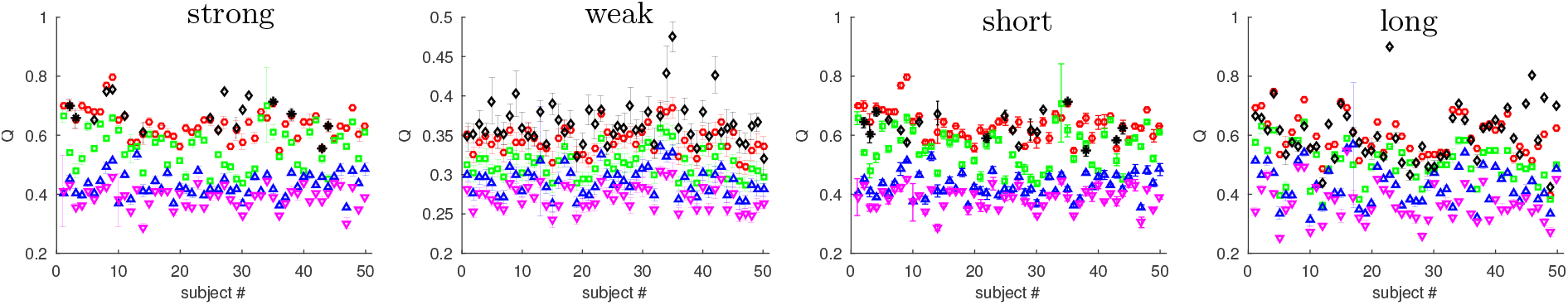
Mean modularity coefficient and their errorbars for the connectome of each subject containing only the long, short, strong or weak links. The most optimal partitions are analyzed for 2, 3, 4 and 5 clusters, and for preset hemispheric division in the cases when not all of the 100 realizations of bicluster partitions correspond to the hemispheric division.

**Figure S3:**
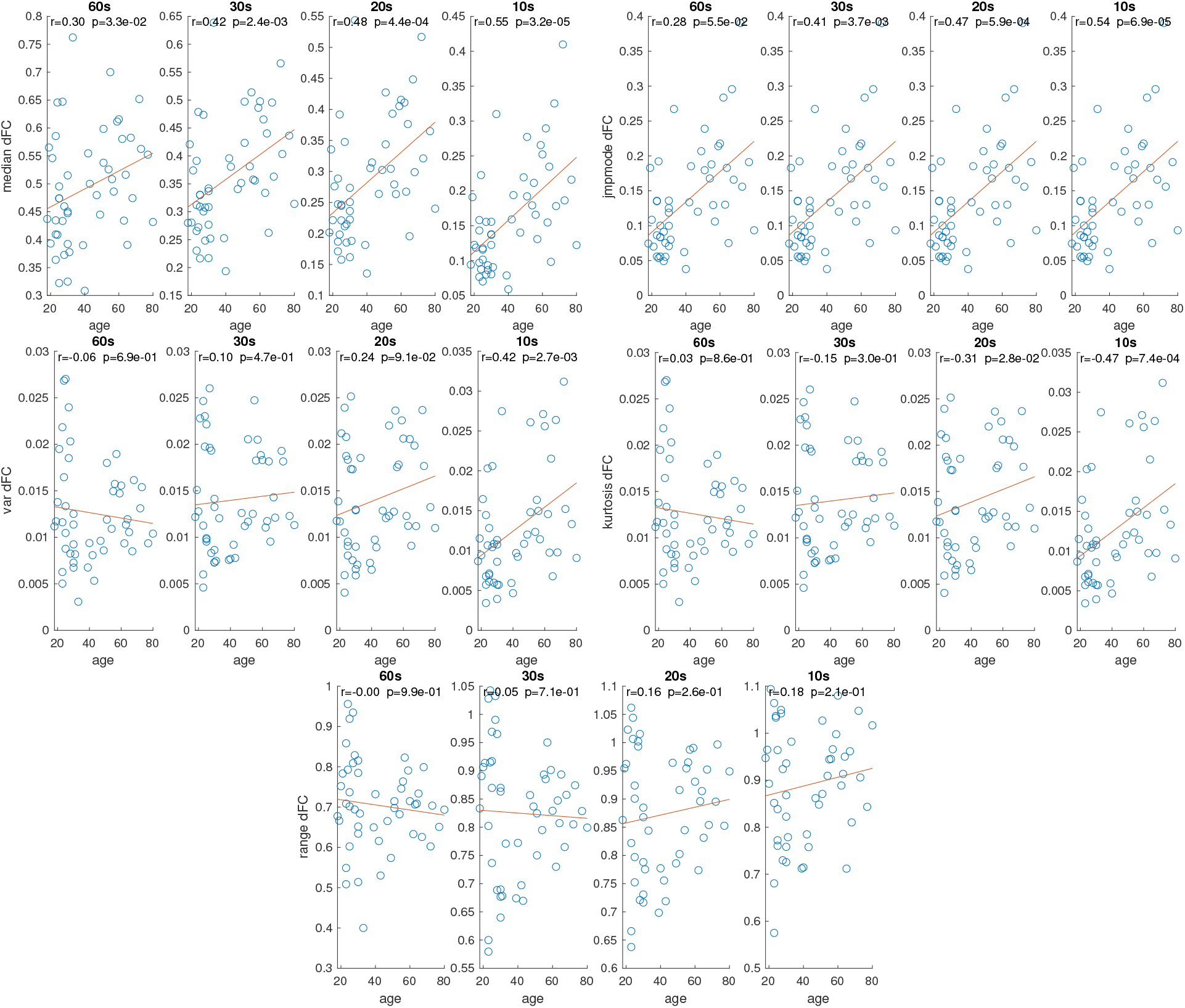
Scatter plots and the correlation of median, mode, variance, kurtosis, and range of dFC, against the age of the subjects for different sizes of the sliding window.

**Figure S4:**
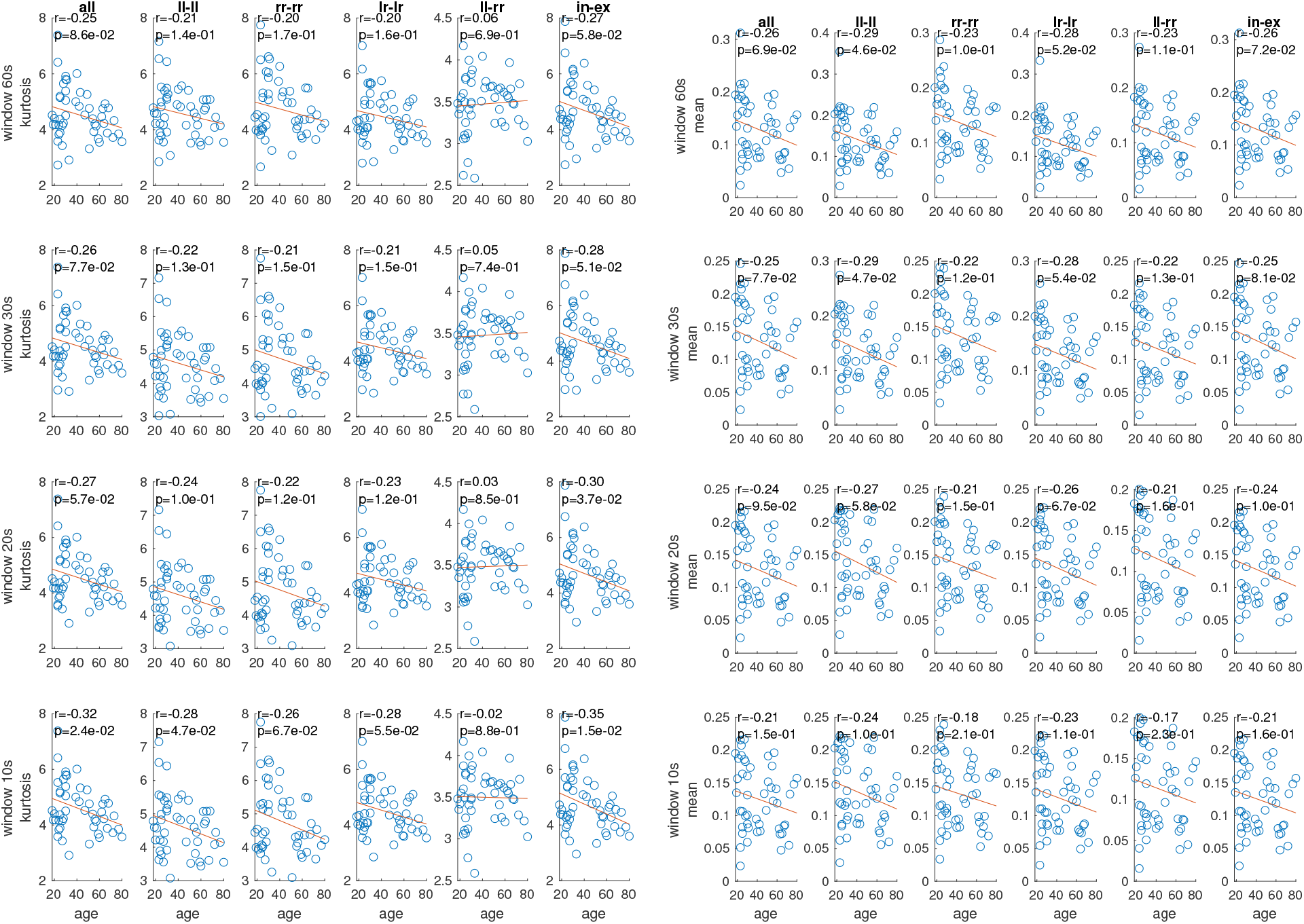
Correlation with age of kurtosis (left) and mean (right) of different spatial components of the empirical MC for different sizes of the sliding window.

**Table S1:** Correlation coefficients with age and their p-values (in brackets) for the number of short and long tracts (shorter and longer than 70*mm*) in and between different lobes. Statistically significant values (*p* < 0.05) are bold, values with *p* < 0.005, *p* < 0.0005 and *p* < 0.00005 are indicated with one, two and three asterisks respectively.

| short ($l < 70mm$) | | | | | | long ($l > 70mm$) | | | | | |
| --- | --- | --- | --- | --- | --- | --- | --- | --- | --- | --- | --- |
|  | Temporal | Cingulate | Frontal | Occipital | Parietal |  | Temporal | Cingulate | Frontal | Occipital | Parietal |
| Temporal | -0.24<br>(8.9e-2) | -0.28<br>(5.1e-2) | -0.09<br>(5.5e-1) | **-0.29**<br>(4.2e-2) | **-0.49\*\***<br>(2.6e-4) | Temporal | -0.08<br>(5.8e-1) | **0.29**<br>(4.0e-2) | 0.09<br>(5.2e-1) | **-0.45\***<br>(1.0e-3) | -0.15<br>(3.0e-1) |
| Cingulate | -0.28<br>(5.1e-2) | 0.22<br>(1.2e-1) | **-0.52\*\***<br>(1.0e-4) | -0.25<br>(8.5e-2) | **-0.49\*\***<br>(3.4e-4) | Cingulate | **0.29**<br>(4.0e-2) | 0.02<br>(8.9e-1) | -0.06<br>(6.8e-1) | -0.25<br>(7.8e-2) | -0.10<br>(5.1e-1) |
| Frontal | -0.09<br>(5.5e-1) | **-0.52\*\***<br>(1.0e-4) | -0.22<br>(1.2e-1) | — | 0.16<br>(2.7e-1) | Frontal | 0.09<br>(5.2e-1) | -0.06<br>(6.8e-1) | **-0.54\*\*\***<br>(4.3e-5) | 0.03<br>(8.2e-1) | -0.12<br>(4.2e-1) |
| Occipital | **-0.29**<br>(4.2e-2) | -0.25<br>(8.5e-2) | — | **-0.50\*\***<br>(2.1e-4) | -0.16<br>(2.5e-1) | Occipital | **-0.45\***<br>(1.0e-3) | -0.25<br>(7.8e-2) | 0.03<br>(8.2e-1) | **-0.42\***<br>(2.7e-3) | **-0.46\***<br>(6.9e-4) |
| Parietal | **-0.49\*\***<br>(2.6e-4) | **-0.49\*\***<br>(3.4e-4) | 0.16<br>(2.7e-1) | -0.16<br>(2.5e-1) | -0.10<br>(4.8e-1) | Parietal | -0.15<br>(3.0e-1) | -0.10<br>(5.1e-1) | -0.12<br>(4.2e-1) | **-0.46\***<br>(6.9e-4) | **-0.57\*\*\***<br>(1.9e-5) |

**Table S2:** Correlation coefficients with age and their p-values (in brackets) for the mean length of short and long tracts (shorter and longer than 70*mm*) in and between different lobes. Statistically significant values (*p* < 0.05) are bold, values with *p* < 0.005, *p* < 0.0005 and *p* < 0.00005 are indicated with one, two and three asterisks respectively.

| short ($l < 70mm$) | | | | | | long ($l > 70mm$) | | | | | |
| --- | --- | --- | --- | --- | --- | --- | --- | --- | --- | --- | --- |
|  | Temporal | Cingulate | Frontal | Occipital | Parietal |  | Temporal | Cingulate | Frontal | Occipital | Parietal |
| Temporal | -0.24<br>(9.4e-2) | 0.27<br>(6.0e-2) | 0.14<br>(3.4e-1) | -0.06<br>(6.9e-1) | -0.06<br>(6.6e-1) | Temporal | **0.47\***<br>**(6.1e-4)** | 0.16<br>(2.6e-1) | 0.05<br>(7.2e-1) | -0.15<br>(3.0e-1) | -0.16<br>(2.6e-1) |
| Cingulate | 0.27<br>(6.0e-2) | 0.03<br>(8.2e-1) | **-0.28**<br>**(4.7e-2)** | -0.03<br>(8.3e-1) | **-0.42\***<br>**(2.6e-3)** | Cingulate | 0.16<br>(2.6e-1) | 0.11<br>(4.4e-1) | 0.08<br>(5.7e-1) | **0.28**<br>**(4.7e-2)** | -0.05<br>(7.4e-1) |
| Frontal | 0.14<br>(3.4e-1) | **-0.28**<br>**(4.7e-2)** | -0.19<br>(1.9e-1) | – | -0.02<br>(8.9e-1) | Frontal | 0.05<br>(7.2e-1) | 0.08<br>(5.7e-1) | **-0.28**<br>**(4.8e-2)** | -0.01<br>(9.4e-1) | **-0.46\***<br>**(8.7e-4)** |
| Occipital | -0.06<br>(6.9e-1) | -0.03<br>(8.3e-1) | – | 0.03<br>(8.5e-1) | 0.03<br>(8.6e-1) | Occipital | -0.15<br>(3.0e-1) | **0.28**<br>**(4.7e-2)** | -0.01<br>(9.4e-1) | -0.18<br>(2.2e-1) | **-0.35**<br>**(1.2e-2)** |
| Parietal | -0.06<br>(6.6e-1) | **-0.42\***<br>**(2.6e-3)** | -0.02<br>(8.9e-1) | 0.03<br>(8.6e-1) | 0.11<br>(4.3e-1) | Parietal | -0.16<br>(2.6e-1) | -0.05<br>(7.4e-1) | **-0.46\***<br>**(8.7e-4)** | **-0.35**<br>**(1.2e-2)** | -0.19<br>(1.8e-1) |

**Table S3:** Correlation coefficients with age and p-values (in brackets) for the mean length of tracts in and between different lobes. Statistically significant values (*p* < 0.05) are bold, values with *p* < 0.005 and *p* < 0.0005 are indicated with one asterisks and two asterisks respectively.

| Lobes |  | Temporal |  | Cingulate |  | Frontal |  | Occipital |  | Parietal |  |
| --- | --- | --- | --- | --- | --- | --- | --- | --- | --- | --- | --- |
|  |  | left | right | left | right | left | right | left | right | left | right |
| Temporal | left | -0.02<br>(8.9e-1) | 0.17<br>(2.3e-1) | **0.31**<br>**(3.1e-2)** | 0.13<br>(3.6e-1) | -0.06<br>(6.6e-1) | -0.11<br>(4.7e-1) | -0.18<br>(2.0e-1) | -0.02<br>(8.9e-1) | 0.24<br>(9.8e-2) | 0.03<br>(8.6e-1) |
|  | right |  | **-0.41\***<br>**(2.8e-3)** | 0.15<br>(2.9e-1) | 0.14<br>(3.2e-1) | 0.27<br>(6.2e-2) | -0.09<br>(5.3e-1) | 0.01<br>(9.5e-1) | -0.02<br>(9.1e-1) | 0.0 (9.8e-1) | 0.02<br>(9.1e-1) |
| Cingulate | left |  |  | 0.09<br>(5.5e-1) | 0.01<br>(9.5e-1) | 0.04<br>(0.78) | **-0.44\***<br>**(1.4e-3)** | -0.26<br>(7.3e-2) | 0.01<br>(9.3e-1) | **-0.38**<br>**(6.3e-3)** | -0.24<br>(8.9e-2) |
|  | right |  |  |  | 0.03<br>(8.3e-1) | **-0.35**<br>**(1.3e-2)** | -0.08<br>(5.6e-1) | 0.08<br>(5.6e-1) | 0.05<br>(7.4e-1) | **-0.29**<br>**(4.1e-2)** | **-0.38**<br>**(6.8e-3)** |
| Frontal | left |  |  |  |  | -0.20<br>(1.7e-1) | **-0.36**<br>**(9.7e-3)** | -0.26<br>(6.7e-2) | **0.30**<br>**3.5e-2** | -0.04<br>(7.8e-1) | 0.01<br>(9.3e-1) |
|  | right |  |  |  |  |  | -0.14<br>(3.3e-1) | -0.07<br>(6.3e-5) | 0.09<br>(5.4e-1) | -0.13<br>(3.7e-1) | 0.10<br>(5.0e-1) |
| Occipital | left |  |  |  |  |  |  | -0.01<br>(9.5e-1) | -0.19<br>(1.8e-1) | 0.13<br>(3.7e-1) | 0.21<br>(1.5e-1) |
|  | right |  |  |  |  |  |  |  | 0.07<br>(6.1e-1) | -0.26<br>(7.4e-2) | -0.04<br>(7.7e-1) |
| Parietal | left |  |  |  |  |  |  |  |  | 0.18<br>(2.1e-1) | **-0.29**<br>**(4.0e-2)** |
|  | right |  |  |  |  |  |  |  |  |  | 0.01<br>(9.3e-1) |

**Table S4:**
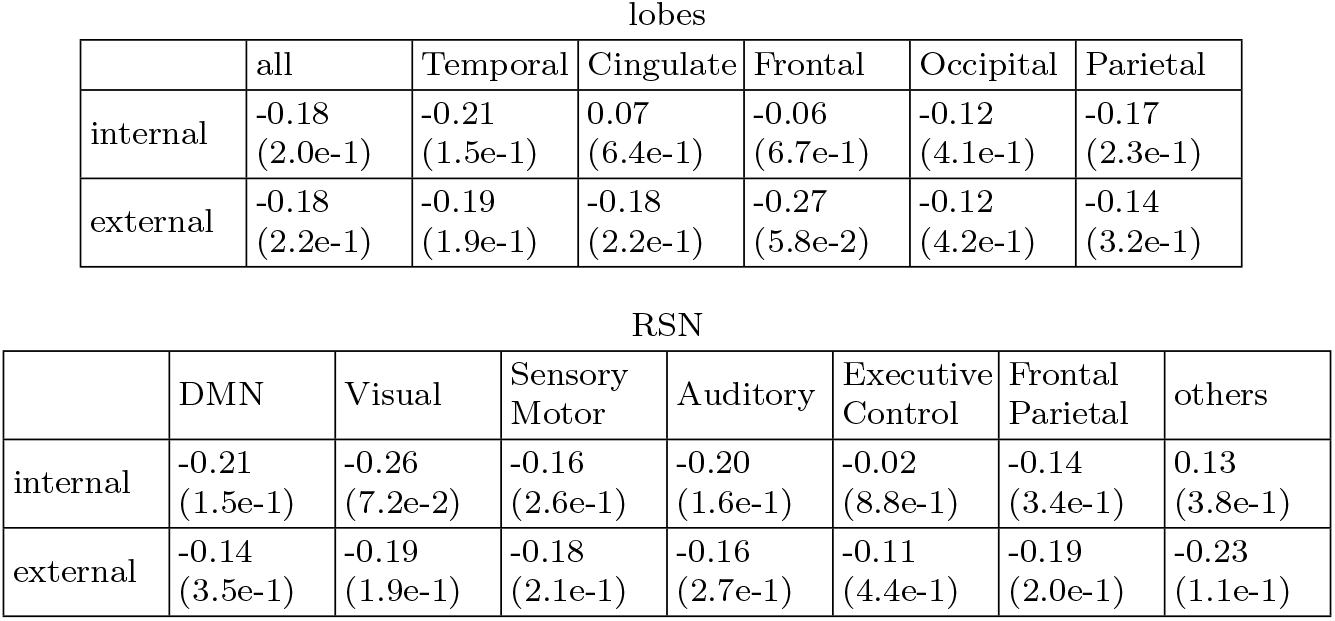
Correlation coefficients with age and p-values (in brackets) for the FC strength of all the internal and external links for a given lobe (top) and RSN (bottom). No statistically significant values (*p* < 0.05) are detected.

| lobes |  |  |  |  |  |  |
| --- | --- | --- | --- | --- | --- | --- |
|  | all | Temporal | Cingulate | Frontal | Occipital | Parietal |
| internal | -0.18<br>(2.0e-1) | -0.21<br>(1.5e-1) | 0.07<br>(6.4e-1) | -0.06<br>(6.7e-1) | -0.12<br>(4.1e-1) | -0.17<br>(2.3e-1) |
| external | -0.18<br>(2.2e-1) | -0.19<br>(1.9e-1) | -0.18<br>(2.2e-1) | -0.27<br>(5.8e-2) | -0.12<br>(4.2e-1) | -0.14<br>(3.2e-1) |

| RSN |  |  |  |  |  |  |  |
| --- | --- | --- | --- | --- | --- | --- | --- |
|  | DMN | Visual | Sensory<br>Motor | Auditory | Executive<br>Control | Frontal<br>Parietal | others |
| internal | -0.21<br>(1.5e-1) | -0.26<br>(7.2e-2) | -0.16<br>(2.6e-1) | -0.20<br>(1.6e-1) | -0.02<br>(8.8e-1) | -0.14<br>(3.4e-1) | 0.13<br>(3.8e-1) |
| external | -0.14<br>(3.5e-1) | -0.19<br>(1.9e-1) | -0.18<br>(2.1e-1) | -0.16<br>(2.7e-1) | -0.11<br>(4.4e-1) | -0.19<br>(2.0e-1) | -0.23<br>(1.1e-1) |

## Notes

### Competing Interest Statement

The authors have declared no competing interest.

